# Spatiotemporal transcriptomic landscape of synovial joint repair – an in vivo murine multimodal model of osteochondral injury

**DOI:** 10.64898/2026.08.05.742782

**Authors:** Rawiya Al Hosni, Francesa Beaton, Andrew Hotchen, Karthik Chary, Nisha Kuzhuppilly Ramakrishnan, Joshua Kaggie, Mark Birch, Andrew McCaskie

## Abstract

**Objective:** The repair response to focal osteochondral injuries frequently fails to truly restore native osteochondral tissue, predisposing the joint to the likelihood of progressive degeneration and post-traumatic osteoarthritis. The biological mechanisms governing the earliest stages of repair in these tissues remain poorly understood, limiting the development of effective regenerative therapies. We therefore aimed to define the early cellular and spatial organisation of repair in a reproducible murine osteochondral injury model by integrating single cell spatial transcriptomics across the whole joint with longitudinal structural imaging and histological analyses.

**Design:** A reproducible, non-critical osteochondral injury was created in the trochlear groove of female C57BL/6 mice. Structural repair was assessed using a multimodal approaching comprising quantitative histology, immunophenotyping, longitudinal magnetic resonance imaging (MRI) and micro-computed tomography (µCT), while whole-joint Xenium spatial transcriptomics at days 3 and 7 defined the cellular and molecular organisation of the early repair response.

**Results:** Spatial transcriptomics demonstrated that the first week after injury is characterised by the emergence of anatomically distinct immune, vascular and stromal microenvironments across the synovial joint. Resolution of the early inflammatory response was accompanied by regional organisation of repair-associated stromal populations by day 7 after injury. The synovium preferentially supported matrix-associated fibro-chondrocyte-like cells, whereas the osteochondral injury itself retained stress-responsive stromal states with comparatively limited representation of matrix-associated populations. These findings indicate that distinct anatomical niches within the joint are associated with transcriptionally distinct stromal cell phenotypes during early repair. Longitudinal MRI and µCT and histological analysis, demonstrated that these early spatial differences in cell phenotype were associated with progressive restoration of osteochondral architecture, with more effective regeneration of subchondral bone and limited restoration of native articular cartilage.

**Conclusions:** This study provides, to our knowledge, the first spatially resolved transcriptomic analysis of the early osteochondral repair response to injury across the whole synovial joint. Our findings demonstrate that the first week after injury establishes spatially organised immune, vascular and stromal cell microenvironments. Furthermore, these data suggest that incomplete cartilage repair may reflect an initial failure to establish and sustain matrix-associated stromal cellular states within the injury niche. These findings identify the early repair microenvironment as a critical determinant of tissue regeneration and provide a rationale for regenerative strategies that target repair with spatial and temporal precision.

## Introduction

Osteochondral injury is an injury to the articular cartilage and the underlying bone of a human joint and is a recognised risk factor for the development of post-traumatic osteoarthritis. Chondral and osteochondral lesions are identified in up to 60% of patients undergoing knee arthroscopy (Curl et al., 1997), yet current treatments do not reliably restore native joint tissues or prevent long-term degeneration. Such degeneration of osteoarthritis (OA) affects more than 500 million people worldwide and is a leading cause of pain and disability, with limited disease-modifying treatment options. Increasing evidence suggests that the biological events occurring during the early response to osteochondral injury play a critical role in shaping subsequent repair quality and long-term joint health, representing an important therapeutic “window of opportunity” for intervention (Mahmoudian et al., 2021). The treatment of early OA, before joint replacement, is an important global area of unmet clinical need.

Despite this importance, the cellular and molecular mechanisms that coordinate these early repair processes remain poorly understood, limiting the development of regenerative and disease-modifying therapies aimed at preserving joint function and preventing subsequent osteoarthritis (Luyten et al., 2012; Madry et al., 2016; Mahmoudian et al., 2021; Lieberthal et al., 2015; Eckstein et al., 2020). Articular cartilage is avascular and relies on diffusion from synovial fluid and subchondral bone for nutrient supply, factors that are thought to limit its intrinsic regenerative capacity following injury (Hunziker, 2002; Armiento et al., 2019; Madry et al., 2021). Despite this, the joint can mount a reparative response under certain conditions. Bone marrow stimulation techniques such as Pridie drilling and microfracture breach the subchondral bone, enabling recruitment of marrow-derived cells and formation of repair tissue (Pridie, 1959; Shapiro et al., 1993; Erggelet & Vavken, 2016). However, the resulting fibrocartilage is mechanically inferior to native hyaline cartilage, with reduced proteoglycan content and disorganised collagen architecture. Consequently, repair tissue often undergoes progressive fibrillation, thinning and loss, contributing to the development of post-traumatic osteoarthritis (Mithoefer et al., 2009; Minas & Ogura, 2016; Solheim et al., 2016). Cell-based therapies, including autologous chondrocyte implantation (ACI) and matrix-associated ACI, aim to improve the quality of repair tissue, but clinical outcomes remain variable and are influenced by patient heterogeneity and procedural differences (Gracitelli et al., 2015; Madry et al., 2021). These limitations are a key gap in our understanding of how the local joint environment regulates early repair responses and determines long-term tissue quality and will be explored in this paper.

Murine models offer powerful tools to investigate the molecular and cellular mechanisms underpinning cartilage repair, enabling genetic manipulation, immunophenotyping, and longitudinal analysis within the same joint (Little & Hunter, 2013). Previous studies have demonstrated that repair capacity is context-dependent, with age, injury type, and local signalling environment influencing outcomes (Kuroda et al., 2006; van der Kraan, 2017). Existing models predominantly demonstrate chronic osteoarthritis, such as destabilisation of the medial meniscus (DMM) and collagenase-induced joint instability, as well as focal chondral injury models that have provided important insights into endogenous cartilage repair (Glasson et al., 2007; Eltawil NM et al., 2009). However, these models either primarily recapitulate progressive joint degeneration or do not fully reproduce disruption of the osteochondral unit and the associated contribution of subchondral bone and bone marrow derived cells to repair. Consequently, the early post injury phase during which immune activation, stromal recruitment, and tissue remodelling are initiated, remains incompletely characterised in vivo. We have developed and validated a reproducible, non-critical murine osteochondral injury model to investigate the early biological events that govern the endogenous response to osteochondral injury. We aimed to establish a multimodal analysis platform integrating quantitative histology, immunophenotyping, non-destructive longitudinal imaging, and spatial transcriptomics to define the structural, cellular and molecular events that shape early repair across the injured joint. Together, this platform provides a framework for understanding the biological processes that underlie the regenerative capacity of articular cartilage and for evaluating strategies designed to enhance this.

## Methods

### Animal housing and husbandry

All procedures were conducted in accordance with the UK Animals (Scientific Procedures) Act 1986 and were approved by the University of Cambridge Animal Welfare and Ethical Review Body (AWERB) under a UK Home Office Project Licence (PP90129760). Mice were housed in ventilated cages in groups of 5 under a 12h dark 12h light cycle and fed a standard diet. Mice underwent 1 week acclimatisation period prior to any procedure. Female wild type C57BL/6 mice received surgeries between 8 and 10 weeks of age. All study plans were reviewed and approved in line with the animal facility prior to any surgeries.

### Development and validation of the murine osteochondral injury model

To establish a reproducible murine model of osteochondral injury suitable for investigating the tissue, cellular and molecular events associated with cartilage repair, a cylindrical osteochondral defect disrupting the articular cartilage, subchondral bone and marrow cavity was created in the patellar (trochlear) groove of female C57BL/6 mice aged 8-10 weeks. The model was developed based on previously published murine osteochondral and chondral injury models (Fitzgerald et al., 2008; Eltawil et al., 2009; Matsuoka et al., 2015) and further refined to maximise surgical reproducibility while minimising damage to the distal femoral epiphysis and growth plate.

During model optimisation, surgical parameters including needle gauge, insertion angle and defect depth were systematically evaluated using cadaveric mouse knees before in vivo experimentation. Hypodermic needles ranging from 21G to 27G were assessed for their ability to generate a consistent cylindrical defect extending through the articular cartilage, subchondral bone and into the marrow cavity. Alternative approaches to standardise defect depth, including plastic depth guards, beads and needle sheaths, were also evaluated but were not adopted as they impaired visualisation of the injury site and increased damage to the surrounding cartilage surface. Based on these optimisation studies, a 26G hypodermic needle inserted to the depth of its bevel was selected for all subsequent experiments, providing consistent access to the marrow cavity while avoiding disruption of the growth plate (Fig. 1).

**Figure 1.**
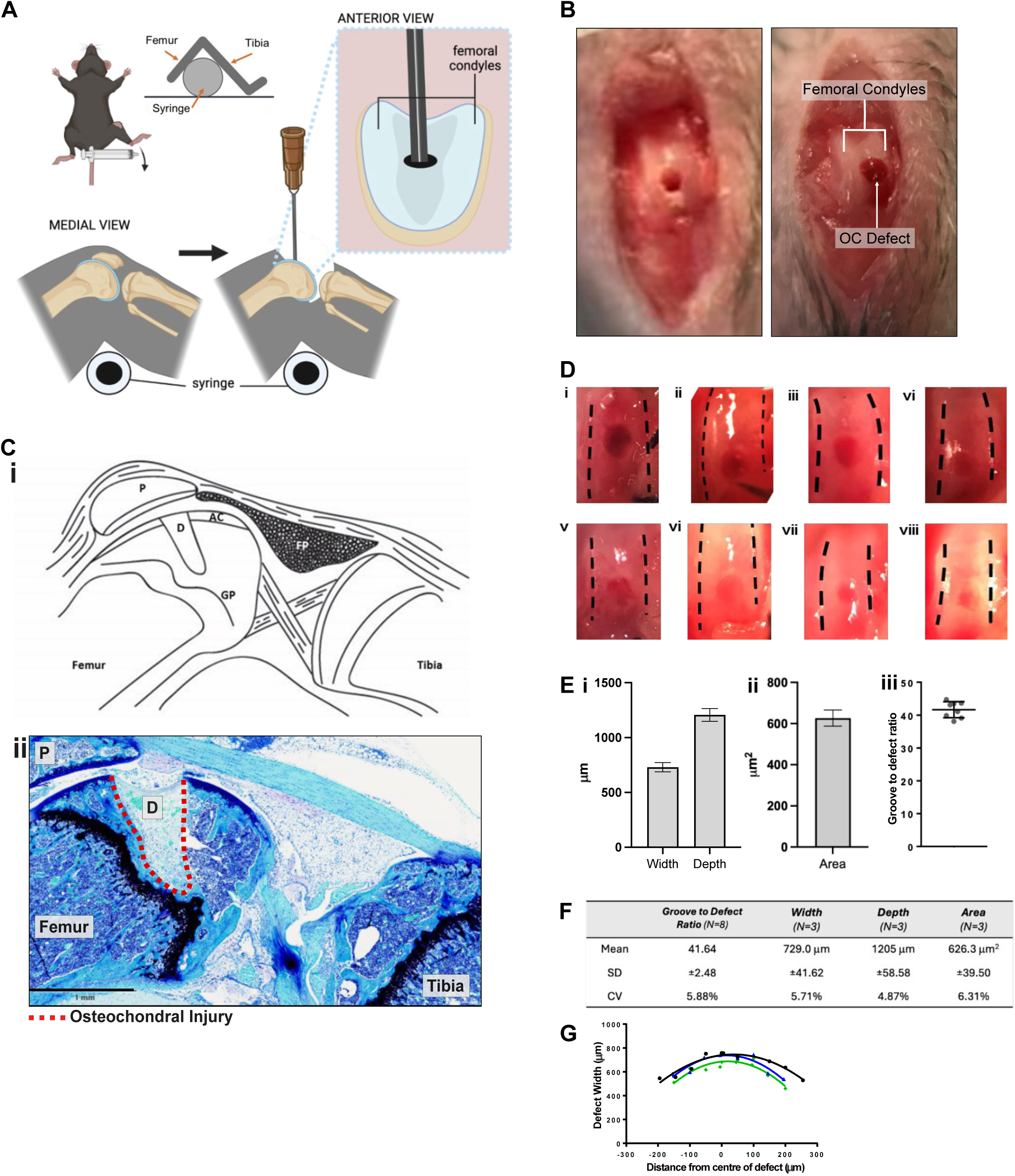
Development and validation of a reproducible osteochondral injury model. (A) Schematic representation of osteochondral injury induction in the murine knee joint, illustrating needle positioning within the femoral condyle in anterior and medial views. (B) Representative macroscopic images of the femoral condyle following injury induction, highlighting the location and morphology of the osteochondral injury. (C) Saggital anatomical schematic of the murine knee joint (i) and corresponding histological section stained with Toluidine blue counterstained with Fast green (ii) confirming defect placement within the femoral condyle, with the injury site indicated (Scale bar= 1 mm). (D) Representative images demonstrating consistency of defect creation across multiple samples. (E) Quantification of defect geometry, including width and depth (i), defect area (ii), and groove-to-defect ratio (N=8) (iii). (F) Summary table of defect measurements showing mean, ± standard deviation (SD), and coefficient of variation. n = 8, 3, 3, 3, defects per group, respectively. (G) Radial profile of defect shape, illustrating defect width relative to distance from the centre of the injury. Data collected from serial sections 50 μm apart. Zero represents the centre of the defect.

General anaesthesia was induced using 4% isoflurane and maintained at 1.5-2% isoflurane in 1-2 L/min oxygen throughout surgery. Mice received pre-operative subcutaneous buprenorphine (1 mg/kg). Following hair removal and skin disinfection, animals were positioned in dorsal recumbency on a heated operating platform beneath a surgical microscope. The left hindlimb was flexed over the barrel of a 2.5 mL syringe and secured to maintain a right angle at the knee. A 0.5-1 cm medial parapatellar skin incision was made using a No. 10A scalpel blade, followed by blunt dissection and incision of the joint capsule adjacent to the patella using a 26G needle. The patella was dislocated laterally to expose the trochlear groove, and the osteochondral defect was created by rotating a 26G hypodermic needle under gentle pressure until the bevel was fully inserted, generating a cylindrical defect extending through the articular cartilage, subchondral bone and into the marrow cavity (Fig. 1A, B). Successful penetration into the marrow space was confirmed intra-operatively by bleeding from the defect.

Following injury creation, the defect was irrigated with sterile saline and a drop of 0.25% bupivacaine applied before the patella was repositioned. The joint capsule was closed using two interrupted 7-0 absorbable sutures (Ethicon), followed by closure of the skin with three to four interrupted 6-0 sutures (Ethicon).

Animals recovered in a warmed incubator and were monitored every 15 minutes during the first postoperative hour until fully ambulatory. Mice were subsequently housed under standard conditions and monitored daily for seven days, including assessment of body weight, wound healing and limb function. Oral meloxicam (1.5 mg/mL; one drop) was administered for 48 hours following surgery. A body weight loss exceeding 15% of pre-operative weight was predefined as a humane endpoint.

To validate the reproducibility of the optimised surgical procedure, osteochondral defects were generated in eight mice and joints were collected 24 hours after injury (Fig. 1D-G). Macroscopic assessment, histological analysis and quantitative histomorphometry were performed to evaluate defect localisation, morphology, width, depth and area. Serial sagittal sections spanning the lesion at ≤50 μm intervals were additionally analysed to assess the consistency of defect geometry throughout the injury.

### MRI and CT scanning

A multi-modal longitudinal imaging protocol was employed to assess the knees at one, four-, and eight-weeks post-injury. This protocol facilitated the evaluation of cartilage morphology, the visualisation of surrounding soft tissue structures, and a three-dimensional assessment of the joint capsule.

Prior to imaging, the mice were initially sedated with 4% isoflurane and subsequently maintained under 1-2% isoflurane during the scan. Their breathing and temperature were continuously monitored using a respiration pneumatic sensor and a rectal temperature probe, ensuring their physiological stability was maintained (breathing rate of 30-50 bpm and temperature between 35-37°C) throughout the scans.

CT data were acquired using a Mediso nanoScan PET/CT system (Mediso Medical Imaging systems, Budapest, Hungary). The murine knees were positioned in a manner consistent with MRI acquisition to ensure reproducible imagining orientation. The scans employed a semicircular single-field-of-view (FOV) acquisition technique with a total of 720 projections. The system operated at a tube voltage of 50 kVp and a current of 980 µA, with an exposure time of 300 milliseconds and a 1:1 binning ratio. The images were reconstructed using the Butterworth method, achieving an isotropic resolution of 22 µm³. The injury was identified by defining a three-dimensional volume of interest over the defect site. Semi-automatic segmentation was performed using Otsu thresholding in VivoQuant (Invicro, Boston, MA) to delineate the injury for three-dimensional visualisation and qualitative assessment.

MRI was performed using a 3T Bruker BioSpec system, utilising an 82 mm circularly polarised volume coil for transmission and a 20 mm surface coil for reception, which was placed over the left hindlimb. To maintain consistency across scans, in vivo imaging of the murine knees was carried out while the limbs were held in a natural, flexed position. Sequence optimisation was conducted on three freshly culled mice. A 3D elliptical phase-encoding scheme was implemented in a FLASH and TurboRARE-based sequence to acquire high-resolution anatomical images (in-plane resolution of 69-78 µm², 500 µm slice thickness, with a total scan time of 22 minutes). This was followed by T2 MSME-based mapping (TR = 1400 ms, 6 echoes with TE1:TE6 = 9.82:58.89 ms, total scan time of 20 minutes) to assess water and collagen content in the cartilage. T2 relaxation values were calculated using Paravision360 v3.3 by manually delineating regions of interest (ROIs) on three slices to demarcate the injury on the femoral condyle and surrounding soft tissue.

### Tissue processing of murine samples for histological analysis

Animals were humanely sacrificed post-operatively by carbon dioxide asphyxiation and tissue samples harvested at various time points between 24 hours and 8 weeks post-surgery. The operated left hind was harvested from all mice. Mice were skinned and limbs dislocated and removed followed by immediate fixation in cold Antigenfix (Diapath, Italy) for 45 mins on ice. The limbs were washed with Phosphate Buffered Saline (PBS) and all tissues underwent 10% formalin fixation for 1hr at room temperature. Muscle tissue surrounding the joint was dissected using a scalpel. The limbs were then decalcified in a 14% Ethylenediaminetetraacetic acid (EDTA) solution at pH 7.2 for 12 days at 4°C under constant rotation with solution changes every 3 days. Samples were paraffin embedded and sagittal sections at 8 µm were cut for histological analysis. Decalcification was assessed by using a needle out of the region of interest to ensure the bone had softened. The femoral notch was identified by the shape of the physeal scar and the presence of cruciate ligaments visible.

### Safranin O staining

Paraffin-embedded tissue sections were stained using Safranin O to assess proteoglycan deposition during osteochondral repair. Sections were deparaffinised in xylene and rehydrated through a graded ethanol series to distilled water. Nuclei were stained using Weigert’s iron haematoxylin for 6 min, followed by rinsing in running tap water for 10 min. Sections were counterstained with 0.05% Fast Green for 3 min, briefly differentiated in 1% acetic acid (10 s), and stained with 0.5% Safranin O for 6 min. Finally, sections were dehydrated through graded ethanol, cleared in xylene, and mounted with a permanent mounting medium.

### Toluidine blue staining

Paraffin-embedded sections were deparaffinized in xylene and rehydrated through graded ethanol to distilled water. Sections were stained with 0.04% Toluidine Blue in 0.1 M sodium acetate buffer for 10 min, rinsed, and counterstained with 0.02% fast green for 3 min. After washing, sections were dehydrated in ethanol, cleared in xylene, and mounted. Toluidine Blue was used to detect acidic tissue components, including glycosaminoglycans.

### Movat’s Pentachrome stain

Paraffin-embedded mouse knee sections were deparaffinized, rehydrated, and stained using the Movat’s Pentachrome method. Elastic fibres were stained with an iron haematoxylin-based solution and differentiated in ferric chloride, followed by sodium thiosulfate treatment. Sections were stained with Alcian Blue (pH 2.5) for mucin, Biebrich Scarlet-Acid Fuchsin for muscle and fibrin, differentiated with phosphotungstic acid, and counterstained with Yellow Stain for collagen. Slides were dehydrated, cleared, and mounted. This method enables simultaneous visualisation of extracellular matrix components and tissue structures.

### Image acquisition

Stained sections were imaged using a Precipoint O8 microscope and digital slide scanner under brightfield illumination. Whole-slide images were acquired at 5x magnification with a resolution of 0.25 µm/pixel, using identical exposure and white balance settings across all samples to ensure quantitative comparability.

### Histomorphometry and scoring

To assess the level of cartilage repair in the pre-clinical osteochondral imaging model, a modified Pineda scoring system was used (Pineda et al., 1992). Histological sections stained with Safranin O / Fast Green / Weigert’s iron haematoxylin were scored blindly by 2 assessors. Where disagreement occurred, sections were reviewed jointly and a consensus score was assigned.

### Immunofluorescence and confocal imaging

Paraffin-embedded tissue sections were deparaffinized in xylene, followed by rehydration through a graded ethanol series. Slides were rinsed in deionized water and equilibrated in PBS.

Antigen retrieval was performed by Proteinase K treatment for 10 minutes at room temperature and subsequently washed in Tris Buffered Saline (TBS) containing 0.025% Triton X-100 with gentle agitation. Permeabilization was performed when targeting intracellular proteins by incubating sections in PBS containing 0.25% Triton X-100 for 10 minutes. Slides were then washed three times in PBS for 5 minutes each. Blocking was carried out using serum from the host species of the secondary antibody (e.g. goat or donkey serum) or with 1% (w/v) bovine serum albumin (BSA) in 0.3% Triton X-100 for 1 hour at room temperature. Primary antibodies were diluted in incubation buffer (1% BSA in TBS or PBS) and applied to sections overnight at 4°C. After washing in PBS, secondary antibodies were applied in the same buffer for 2.5 hours at room temperature in the dark. Slides were washed again in PBS, followed by nuclear counterstaining with DAPI. Sections were rinsed in PBS and mounted using anti-fade mounting medium. Images were taken using the Leica SP8 confocal microscope.

### Immunohistochemistry staining

After rehydration to water, samples were treated to unmask antigens through an antigen retrieval procedure. For the identification of collagen proteins, samples were incubated at 37°C in 0.1% hyaluronidase (Sigma) for 50 minutes before washing in PBS for 5 minutes and continuing to the blocking step. Other samples were incubated in 10 mM sodium citrate (Sigma) buffer adjusted to pH 6 containing 0.05% Tween (Sigma). This method was optimised and a final procedure of incubating slides in the buffer at 78°C to 83°C was utilised to help prevent tissue detachment from the slides. After this, slides were left under cold running tap water for 10 min and then blocked.

Slides were blocked for 90 minutes at room temperature in 10% goat serum (Vector Laboratories) diluted in blocking buffer; PBS containing 0.1% Tween20 (Sigma) (PBS-T). Primary antibodies were diluted in blocking buffer at concentrations indicated in Table 3 and applied to sections before incubation overnight at 4°C.

The following day slides were washed in PBS four times for 5 minutes each before incubation in 3% (v/v) hydrogen peroxide (Sigma) for 15 minutes. All incubation steps on the second day were performed at room temperature. Slides were washed three times the host species of the primary antibody used, either an anti-rabbit kit (Rabbit Specific HRP/DAB (ABC) Detection IHC kit, Abcam) was used or an HRP-conjugated secondary antibody (see table).

When using the anti-rabbit kit the manufacturer’s instructions were followed. Briefly, samples were incubated for 10 minutes with biotinylated goat anti-rabbit, washed four times in PBS (5 minutes each) and incubated with the streptavidin peroxidase solution for 10 minutes. Otherwise, when using an HRP-conjugated secondary, the antibody was diluted in 1% BSA and slides incubated for 1 hour.

After secondary antibody incubation or the streptavidin peroxidase incubation (for the kit samples) slides were washed four times in TBS, 5 minutes each. Samples were then incubated with DAB substrate for the time quoted in Table 2 before stopping the enzyme reaction by submersion of the slide in PBS. Finally, samples were counterstained with methyl green for 5 minutes at 60°C before rinsing in tap water followed by 6 to 8 dips in 0.05% acetic acid before dehydration and mounting in DPX solution.

**Table 1.** Modified Pineda cartilage epair scoring.

|  |  |
| --- | --- |
| ***Filling of defect area (including bone & bone marrow)*** |  |
| 125% | 1 |
| 100% | 0 |
| 75% | 1 |
| 50% | 2 |
| 25% | 3 |
| 0% | 4 |
| ***Reconstruction of osteochondral junction*** |  |
| Yes | 0 |
| Almost | 1 |
| Not close | 2 |
| ***Matrix staining (bone & cartilage)*** |  |
| Normal | 0 |
| Reduced staining | 1 |
| Significantly reduced staining | 2 |
| Faint staining | 3 |
| No stain | 4 |
| ***Cell morphology*** |  |
| Normal Reduced staining | 0 |
| Most hyaline and fibrocartilage | 1 |
| Mostly fibrocartilage | 2 |
| Some fibrocartilage, but mostly nonchondrocytic cells | 3 |
| Nonchondrocytic cells only | 4 |

**Table 2.** Primary and secondary antibodies used for immunofluorescence and immunohistochemistry.

| Antigen | Catalog No. | Species | Company | Dilution | DAB incubation | IF incubation |
| --- | --- | --- | --- | --- | --- | --- |
| **Primary antibodies** |  |  |  |  |  |  |
| *Type I Collagen* | ab21286 | Rabbit pAb | Abcam | 1 in 100 | - | Overnight |
| *Type II Collagen* | ab34712 | Rabbit pAb | Abcam | 1 in 400 | - | Overnight |
| *Type X Collagen* | 234196 | Rat mAb | Abcam | 1 in 100 | - | Overnight |
| *Lubricin* | Ab28484 | Mouse mAb | Abcam | 1 in 150 | - | Overnight |
| *NIMP* | ab2557 | Rat mAb | Abcam | 1 in 100 | 6 minutes | - |
| *CD68* | ab125212 | Rabbit pAb | Abcam | 1 in 100 | 3 minutes | - |
| **Secondary antibodies** |  |  |  |  |  |  |
| *HRP-conjugated anti-rat* | ab205720 | Goat | Abcam | 1 in 2,500 |  |  |
| *Anti-Rabbit (AF488)* | ab150085 | Goat | Abcam | 1 in 500 |  | 2.5 Hrs |
| *Anti-Rabbit (AF555)* | ab1500678 | Goat | Abcam | 1 in 500 |  | 2.5 Hrs |

### Quantification of vascular and synovial parameters

#### Tissue preparation and staining

Paraffin sections containing the periarticular and appositional synovial regions to the cartilage injury and adjacent repair cartilage were stained with toluidine blue (See Toluidine blue staining protocol) to visualise tissue architecture and vascular structures. Toluidine blue provides strong contrast between cartilage matrix, synovium, and erythrocyte-filled lumina, allowing consistent delineation of blood vessels without additional immunostaining.

For each time point, one section per mouse was analysed (n = 3 mice per time point). Regions of interest encompassing the synovial tissue and adjacent repair cartilage were selected for morphometric analysis. All image selection and subsequent quantification were performed blinded to experimental group.

#### Quantification of vessel area fraction in periarticular and appositional synovium

Toluidine blue-stained sections were used to quantify vascularity within discrete periarticular and appositional synovial regions relative to the defect. Digital images were analysed using ImageJ/FIJI (v2.9). For each section, two regions of interest (ROIs) were manually delineated based on histologic landmarks: The periarticular synovium was defined as the synovial lining contiguous with the infrapatellar (fat pad-associated) synovial fold, characterized by a thin intimal layer, underlying loose connective tissue, and proximity to adipose tissue; the appositional synovium was defined as the synovial lining directly overlying or bordering the cartilage defect, typically forming a denser, more cellular layer juxtaposed to the repair surface.

Within each ROI, vessel cross-sections were identified morphologically as round or oval lumina devoid of toluidine staining and bordered by basophilic endothelial rims. The selected ROI was duplicated (Image → Duplicate selected region only), converted to 8-bit grayscale, and thresholded to isolate bright luminal spaces (Image → Adjust → Threshold, Otsu or Default algorithm). The upper threshold slider was manually adjusted to retain only clear vessel lumina. Segmentation was performed using Analyse Particles (size filter: 20–∞ px²; Exclude on edges enabled). The vessel area fraction (VAF) for each region was calculated as:

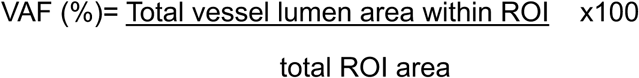

Objects smaller than 20 µm² or larger than 2000 µm² were excluded to eliminate debris and merged lumina. One toluidine blue–stained section per mouse was analysed (n = 3 mice per time point), with both periarticular and appositional ROIs quantified independently.

#### Quantification of vessel size

Vessel size was determined from toluidine blue–stained sections by measuring the equivalent circular diameter (Dₑq) of individual vascular lumina identified within periarticular and appositional synovial regions. Segmentation and morphometric analysis were performed in ImageJ/FIJI using the *Analyze Particles* function. Mean vessel diameters were calculated for each region and time point (1, 4, and 8 weeks; *n* = 3 mice per time point; 1 section per mouse).

#### Measurement of synovial thickness

Synovial thickness was quantified on toluidine blue–stained sections using ImageJ/FIJI (v2.9) with the BoneJ plugin. The synovial ROI was manually delineated and converted to an 8-bit binary mask (white = synovium). Local thickness was computed using *BoneJ → Particle Analyser* (2-D mode, calibrated in µm), and results were expressed as the area-weighted mean thickness ± SD per ROI. Analyses were performed on one section per mouse (*n* = 3 per time point), separately for periarticular (fat pad–adjacent) and appositional (defect-overlying) synovium.

### Spatial transcriptomics

Four mice were culled at 3 (n=7) and 7 days (n=2) following an osteochondral injury. The hind limbs were fixed in 10% formalin for 24hrs. The samples were carefully trimmed to remove excess muscle and soft tissues around the bone and then decalcified in 0.5 M EDTA (pH 8.0) prepared using nuclease-free deionised water. Samples were fully submerged in decalcification solution at a minimum ratio of 20:1 solution volume to tissue volume and maintained under gentle agitation throughout the procedure. Decalcification was performed for 3 days at 4°C, with solution replacement every 24 h. Decalcification was confirmed by manual assessment of bone bendability and piercing the bone using a needle. Tissues were washed 3× in 1× PBS (pH 7.4) to remove residual EDTA, processed through graded alcohols, and embedded in paraffin wax using a routine histology workflow. Paraffin blocks were trimmed to contain the ROI and sectioned at 5 µm thickness for subsequent placement onto a Xenium slide. One representative section containing the centre of the osteochondral defect was selected from each animal for Xenium analysis. The slides were subsequently heated to 60°C for 3 hrs and dried in a desiccator for 2 hrs. Slides were then processed using the xenium workflow protocol. Sections from each sample were reserved for RNA quality assessment using a TapeStation. RNA integrity was evaluated based on fragment size distribution and samples with DV200 values ≥200 bp were considered suitable for the Xenium workflow.

### Data processing and integration

Xenium raw outputs were processed using Xenium Explorer (v3.1) for cell segmentation, transcript assignment, and quality control. Downstream analyses were performed in Seurat (v5.0, R). Cells with <200 detected genes or >15% mitochondrial transcripts were removed, and expression counts were normalized to total transcripts per cell and log transformed.

To directly compare cellular states across time, Day 3 and Day 7 datasets were integrated into a shared reference space using Seurat’s reciprocal principal-component analysis (PCA) pipeline. This approach identifies shared cellular states across datasets to align shared transcriptional states while minimising technical and temporal effects.

The integrated object was then subjected to PCA. An elbow plot was used to determine the number of informative principal components (PCs) to retain. The variance curve began to plateau between PC6 and PC8, indicating diminishing returns in explained variance; therefore, the first 10 PCs were selected for downstream UMAP embedding and Leiden clustering.

After integration and dimensionality reduction, data were visualized in two dimensions using Uniform Manifold Approximation and Projection (UMAP), and clustering was performed on the integrated assay using the Leiden algorithm (resolution = 0.6) to define transcriptionally distinct cell populations. The Leiden clustering resolution parameter was set to 0.6, which provided the best balance between cluster granularity and biological interpretability. This value yielded stable clusters consistent with known joint cell populations and marker gene expression patterns.

Cell-type identities were assigned based on the top 10 differentially expressed genes per cluster (Wilcoxon rank-sum test, adjusted *p*< 0.05) and validated against canonical marker genes from the Xenium Mouse Tissue Atlas panel (379 genes).

### Cell-type assignment and visualization in Xenium Explorer

Cell identities determined in Seurat were exported as metadata and mapped back to the spatial dataset using the Xenium Explorer annotation import function. Cell cluster labels were merged with the Xenium cell-by-feature matrix via unique cell barcodes. These annotations were then visualised as cell types within Xenium Explorer (v3.1) to assess spatial localisation of each population across the joint tissues at Day 3 and Day 7.

### Functional characterisation and spatial mapping of fibro-chondrocyte-like stromal cell states

Gene Ontology (GO) over-representation analysis was performed using the PANTHER Classification System (GO Biological Process, complete) with the *Mus musculus* genome as reference. Enrichment significance was assessed using Fisher’s exact test with Benjamini-Hochberg false discovery rate (FDR) correction, and terms with FDR< 0.05 were retained. Enriched GO terms were manually curated and grouped into biologically relevant functional states reflecting fibro-chondrocyte-like stromal cell (FCSC) behaviour, including matrix-producing, activated fibroblast-like, tissue remodelling, and stress/injury-responsive states. Gene sets for each functional state were generated from the union of genes contributing to GO terms within each category and used for downstream analysis.

Spatial transcriptomic data from Xenium Day 3 and Day 7 samples were analysed in R using Seurat (v4+). Pre-processed Seurat objects containing Xenium gene expression counts were annotated using cell-type labels exported from Xenium Explorer, and analyses were restricted to cells classified as fibro-chondrocyte-like stromal cells (FCSCs). Data were log-normalised using Seurat’s default LogNormalize method (scale factor = 10,000). Functional state activity was quantified at the single-cell level using Seurat’s AddModuleScore function (ctrl = 10, nbins = 10), generating relative enrichment scores for each gene set by comparing average expression of signature genes to matched background gene sets. For each cell, a dominant functional state was assigned based on the highest module score across the four programs.

To account for transcriptional ambiguity, a confidence metric was calculated as the difference between the top and second-highest module scores. Cells with confidence values <0.10 were classified as Mixed/Uncertain. To exclude low-quality or misannotated cells, a core FCSC identity score was defined and cells below the 5th percentile of this distribution were reassigned to a Low Core/Off Target category. Anchor marker genes for each functional state (e.g., *Mfap4*, *Mfap5*, *Htra3*, *Fxyd1*) were selected based on specificity and biological relevance and used for validation and visualisation. Functional states were mapped back onto spatial coordinates to assess their distribution across anatomical regions, including the injury site, appositional synovium, and periarticular synovium, at day 3 and day 7 post-injury.

### Quantification and statistical analysis

Data are presented as mean ± SD unless otherwise stated. Statistical analyses were performed using GraphPad Prism v10. For comparisons involving two independent groups, unpaired two-tailed Student’s *t*-tests with Welch’s correction were used where appropriate. For experiments involving multiple groups or variables, one-way or two-way analysis of variance (ANOVA) was applied as indicated, followed by Uncorrected Fisher’s LSD multiple comparisons testing.

For analyses of synovial and vascular parameters across time and anatomical regions, ordinary two-way ANOVA was used to assess the effects of time, region, and their interaction. Where appropriate, individual data points represent biological replicates. Sample sizes (n) refer to the number of independent animals per condition, as specified in the corresponding figure legends.

Statistical significance is indicated as follows: *P* < 0.05*, <0.01, and <0.001*. Exact statistical tests and sample sizes for each analysis are detailed in the corresponding figure legends.

## Results

### Histological characterisation of osteochondral repair following surgical injury

Histological evaluation of joints collected 24 hours, 1, 2, 4 and 8 weeks following osteochondral injury demonstrated progressive structural remodelling of the osteochondral unit throughout the repair process (Fig. 2A). During the first two weeks after injury, the defect margins remained well defined, clearly demarcating the injured cartilage from the adjacent intact articular surface. By 4 weeks, repair tissue increasingly bridged the defect and the distinction between native and repair tissue became less apparent, coinciding with progressive subchondral bone formation. At 8 weeks, the osteochondral architecture was largely restored, with re-establishment of the cartilage-bone interface and formation of a proteoglycan-containing cartilage-like surface. Within the deeper aspect of the defect, progressive trabecular bone formation was accompanied by reconstruction of the marrow cavity and integration of the repair tissue with the surrounding bone.

**Figure 2.**
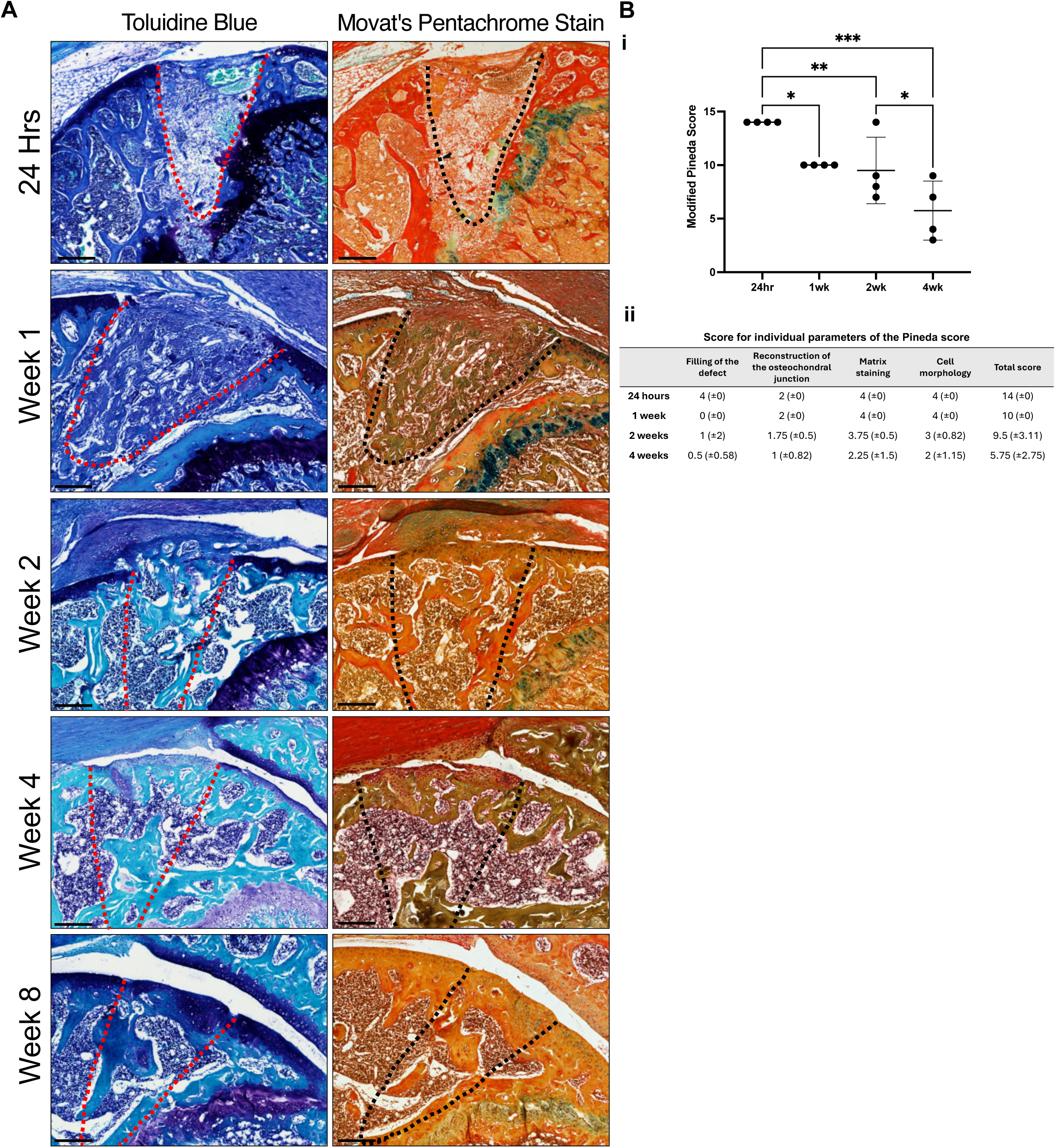
Temporal changes in matrix composition during osteochondral repair. (A) Histological representation of osteochondral injuries at 24hrs, 1-, 2-, 4-, and 8- weeks following injury. Tissue is stained with Toluidine Blue/ Fast Green. Sections are also stained with Movat’s Pentachrome stain, staining for Elastic fibres (black to blue/black); Nuclei (blue/black); Collagen (yellow); Reticular fibres (yellow); Mucin (bright blue); Fibrin (bright red); Muscle (red). Scale bar= 200 μm (B) Articular cartilage repair Pineda Score 24hrs, 1-, 2-, and 4-weeks following injury. Scores derived from histological sections stained with Safranin O/ Haematoxylin (n= 4) (i), summary of individual parameters used to assign Pineda score. Data represented as mean ± SD (ii). Statistical analysis performed using one-way analysis of variance, uncorrected Fisher’s least significant difference; ***p < 0.001; **p < 0.01; *p < 0.05; n= 4).

Analysis of cellular and extracellular matrix composition revealed distinct temporal changes throughout repair. At 24 hours, the defect was predominantly occupied by a provisional clot containing abundant erythrocytes and scattered mononuclear cells. By 1 week, the superficial repair tissue had condensed into a dense cellular layer orientated parallel to the articular surface, while the underlying defect remained highly cellular with focal regions of early proteoglycan deposition identified by Movat’s Pentachrome staining. From 2 weeks onwards, repair became increasingly spatially organised. Discrete regions of proteoglycan-rich matrix accumulated at the articular surface, consistent with early cartilage repair, while the deeper defect underwent progressive remodelling through formation of interconnected subchondral and trabecular bone, supporting restoration of the marrow compartment. Although repair tissue continued to mature throughout the study, the regenerated cartilage remained morphologically distinct from the adjacent native articular cartilage at 8 weeks.

To quantitatively assess repair progression, histological sections through the centre of each defect were independently scored by two blinded observers using the modified Pineda scoring system (Fig. 2B). Histological scores improved progressively over time, reflecting increased defect filling, restoration of the osteochondral junction, improved matrix staining and cellular morphology. Nevertheless, variability between animals remained evident, and complete restoration of native cartilage architecture was not achieved within the experimental period.

### Extracellular matrix deposition after osteochondral injury

To characterise tissue composition and organisation at later stages following an osteochondral injury, immunofluorescence staining was performed for key skeletal lineage matrix markers at 8 weeks (Fig. 3). Strong COL1A1 signal was detected throughout the defect region, and co-localised with the boundary of the native trabecular bone. COL2A1 expression was limited to a narrow layer at the defect surface and contiguous with adjacent native articular cartilage, suggesting restricted hyaline cartilage regeneration. COL10A1 was enriched along the tide mark between the repair tissue and subchondral bone, further supporting hypertrophic chondrocyte activity. Lubricin (PRG4) was detected along the surface of the repaired tissue, demonstrating re-establishment of a lubricating boundary layer that included the restored cartilage.

**Figure 3.**
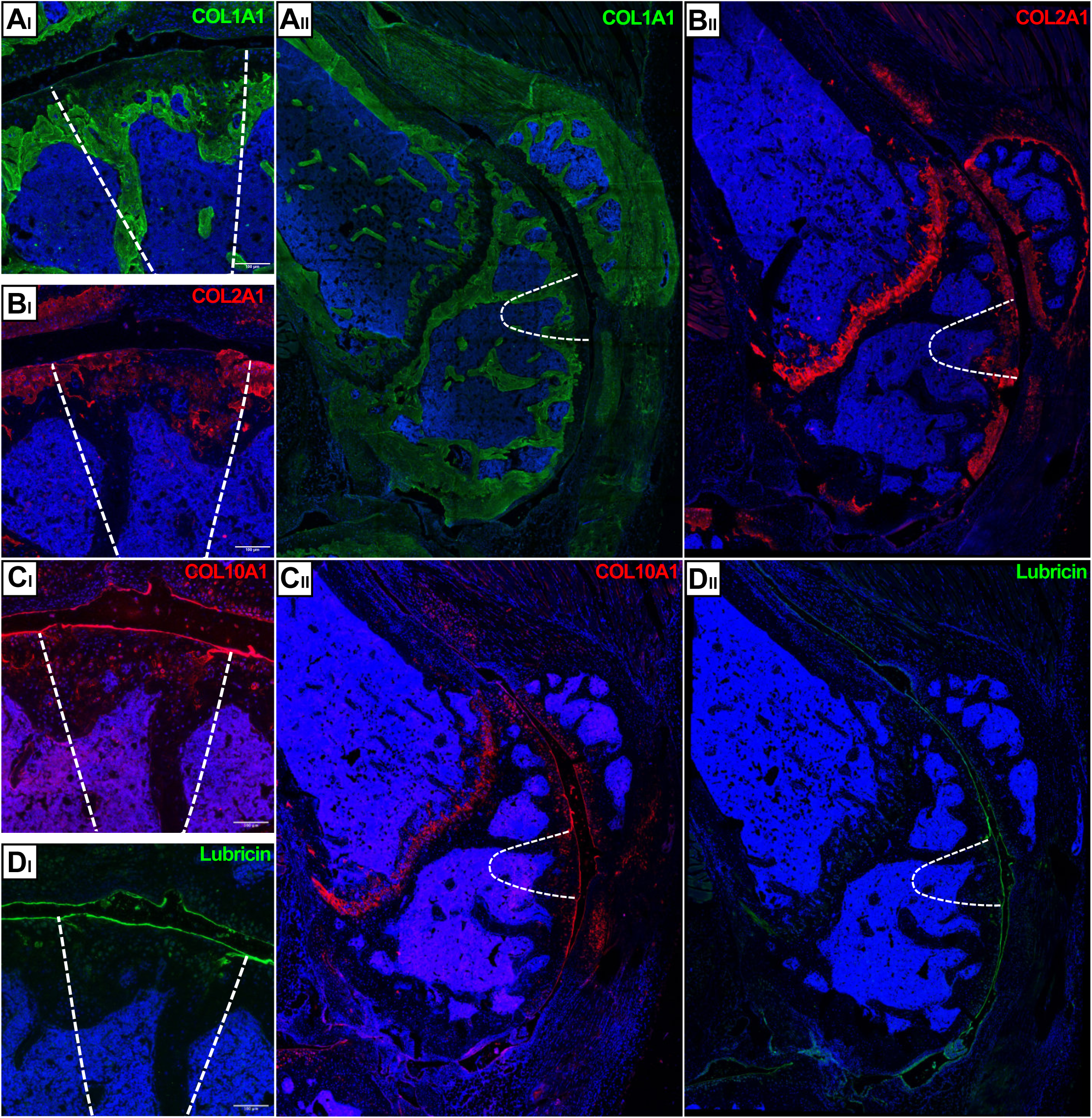
Matrix marker expression in repair tissue at 8 weeks post-osteochondral injury. Immunofluorescence images matrix associated markers at 8- weeks post osteochondral injury. (A) represents COL1A1; (B) COL2A1; (C) COL10A1; (D) Lubricin staining of injury (I) and femoral epiphysis (II). Injury highlighted with white dotted line. Scale bar= 100 μm.

### Longitudinal micro-CT imaging enables non-destructive assessment of osteochondral repair

To follow the progression of bone remodelling within the osteochondral defect in vivo, longitudinal micro-computed topography (μCT) imaging was performed at 1-, 4- and 8-weeks following injury (Fig. 4). At week 1, the defect was clearly visible with distinct loss of subchondral bone, evidenced by a loss of mineralised tissue at the injury site. By week 4, new mineralised tissue was observed, with new bone bridging at the periphery of the injury. By week 8, the defect demonstrated progressive infilling with mineralised tissue occupying much of the defect space and integration with existing subchondral bone, although the architecture remained irregular compared with the surrounding intact bone. These μCT observations confirm a progressive restoration of mineralised tissue within the defect between 1- (n=4) and 8-weeks (n=4) post-injury.

**Figure 4.**
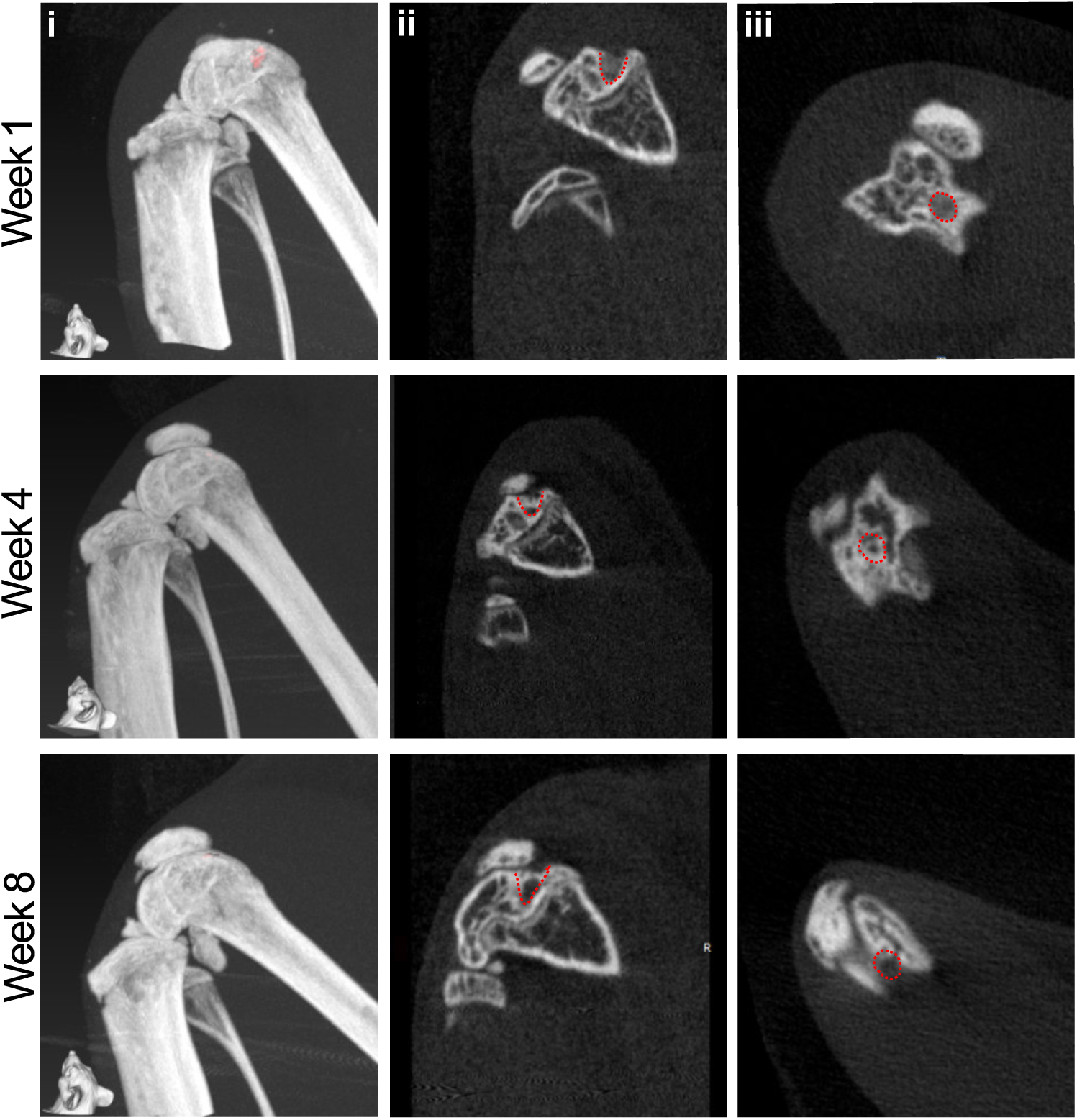
Longitudinal μ-CT assessment of osteochondral repair. Representative image of longitudinal m-CT of a single mouse conducted at 1-, 4- and 8- weeks following injury. 3D representation of the knee with the injury highlighted with a solid red on the femoral condyle (i), sagittal view of the injury highlighted in a red dashed line (ii), and coronal view of the injury highlighted with a red dashed line (iii).

### Longitudinal MRI imaging and histology reveal dynamic synovial and cartilage changes

To characterise soft tissue changes following osteochondral injury, longitudinal MRI was combined with histological analysis and quantification of synovial vascularity (Fig. 5). Representative MRI scans acquired at weeks 1- (Fig. 5A-C), 4- (Fig. 5D-F) and 8-weeks (Fig. 5G-I) post-injury showed hyperintensity near the injury site and synovial thickening at week 1, consistent with early post-injury oedema and synovial activation. Signal intensity and synovial tissue thickening increase at week 4 (Fig. 5D-E), partially resolving by week 8 (Fig. 5G-H), indicating dynamic changes of synovial tissue during repair.

**Figure 5.**
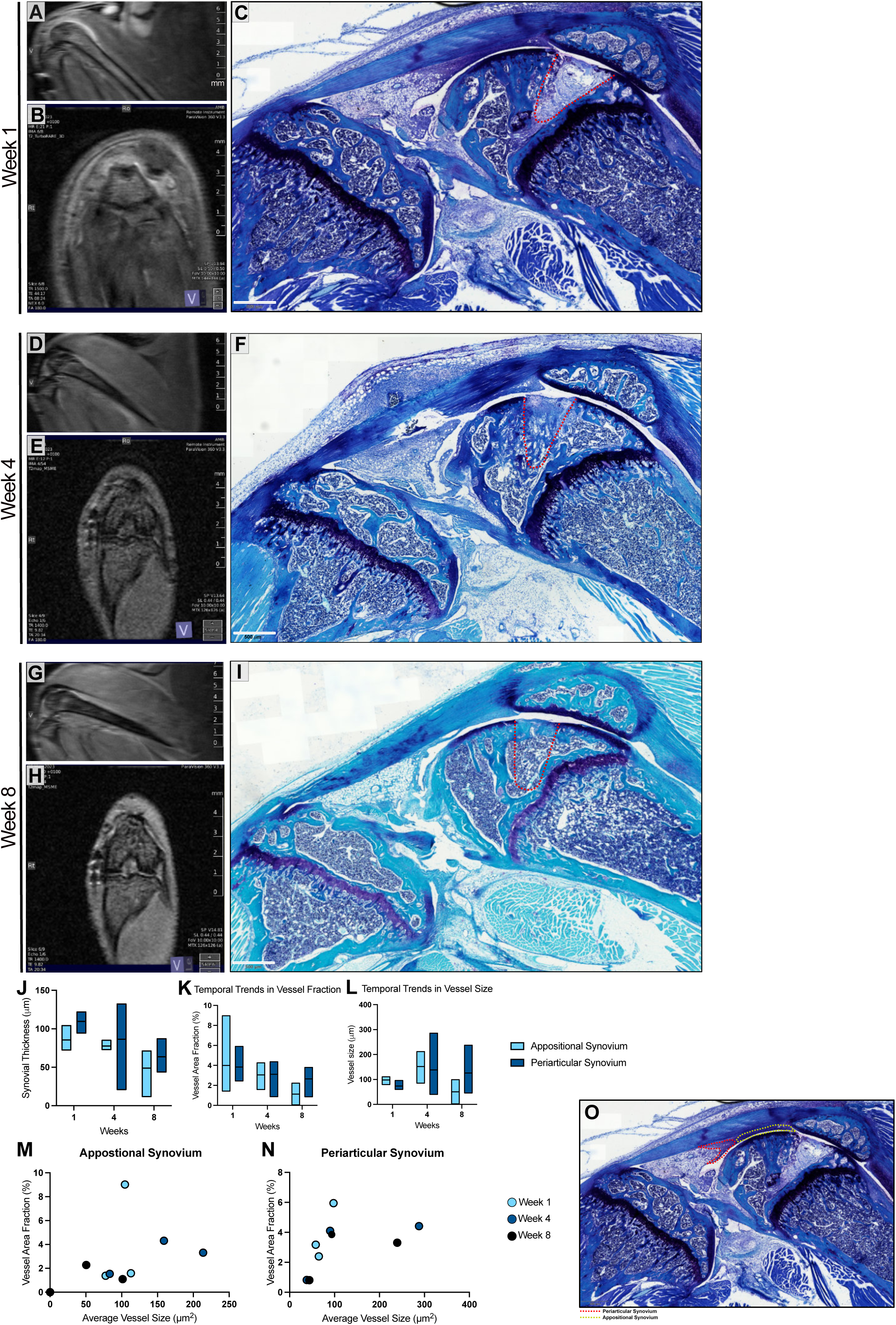
Temporal synovial changes following an osteochondral injury: MRI, histology, and vessel quantification. (A-B, D-E, G-H) Representative MRI images of the murine knee joint acquired longitudinally from the same animal at 1-, 4-, and 8-weeks post-injury, shown in sagittal (A, D, G) and coronal (B, E, H) views, demonstrating temporal changes in intra-articular signal intensity consistent with evolving synovial and joint responses. (C, F, I) Representative Toluidine Blue-stained histological sections from separate animals at matched timepoints, illustrating the osteochondral defect (outlined with red dashed line) and progressive changes in cartilage and subchondral bone architecture. MRI images are from a single representative mouse followed over time; histological sections are from independent animals at each timepoint (n= 4, 6, 4, respectively per timepoint at 1-, 4-, 8- weeks). Changes in synovial architecture were measured using histological sections stained with Toluidine blue. (J) Quantification of synovial thickness within appositional and periarticular synovial regions at 1-, 4-, and 8-weeks post- injury. (K) Vessel area fraction (%) across timepoints, demonstrating changes in vascular density within synovial compartments. (L) Average vessel size (µm) measured over time in appositional and periarticular synovium. (M-N) Relationship between vessel size and vessel area fraction within appositional (M) and periarticular (N) synovium at each timepoint, with individual data points representing biological replicates. (O) Representative Toluidine Blue- stained histological section illustrating anatomical regions used for quantification, including appositional synovium (yellow dashed line) and periarticular synovium (red dashed line). Data are presented as mean ± SD where applicable. n= 4, 6 and 4 animals per timepoint (1-, 4-, 8- weeks, respectively). Statistical analysis was performed using an ordinary two-way ANOVA. Scale bars: 500 µm.

To correlate MRI changes with the synovial tissue architecture, histology and vascular structure quantification were performed on tissue from the same timepoints (Fig. 5C, F, I). Toluidine blue staining distinguished between the appositional synovium directly overlying the defect and the periarticular synovium within the infrapatellar fat pad (Fig. 5O). At week 1, the appositional synovium appeared thickened with visible vascular structures. Thickness and vascularisation increased further at week 4 and was almost completely resolved by week 8 (Fig. 5I).

To further assess these temporal changes quantitatively, synovial thickness was measured in both regions (week 1 (n=3); week 4 (n=5); week 8 (n=4)) (Fig. 5J). The appositional synovium remained consistently thicker than the periarticular lining at all timepoints, with maximal expansion observed between weeks 1-4. Vessel area fraction decreased progressively over the 8-week period (Fig. 5K), while vessel size peaked at week 4 in the appositional region before declining at week 8 (Fig. 5L), indicating a transient vascular enlargement during early repair. Average vessel size compared to vessel area fraction (Fig. 5M-N) demonstrated that an increase in synovial vascularity was associated with expansion of vascular area and dilation of individual vessels, particularly in the appositional synovium.

These findings demonstrate a pronounced reversible inflammatory and vascular response within the appositional synovium following injury, most evident in appositional regions in direct contact with the defect.

### Spatial transcriptomics reveals dynamic immune and stromal cell populations in the joint during early osteochondral repair

To further characterise early cellular responses following an osteochondral injury, Xenium in situ spatial transcriptomics was performed on mouse knee joints at day 3 and day 7 post-injury (Fig. 6). Sections spanning the osteochondral defect, appositional and periarticular synovium, and adjacent joint tissues were processed using the Mouse Tissue Atlas 379-gene panel, followed by in situ sequencing and high-resolution imaging to generate spatially resolved gene expression maps across the entire joint to enable mapping of immune, vascular, and stromal populations during early osteochondral repair.

**Figure 6.**
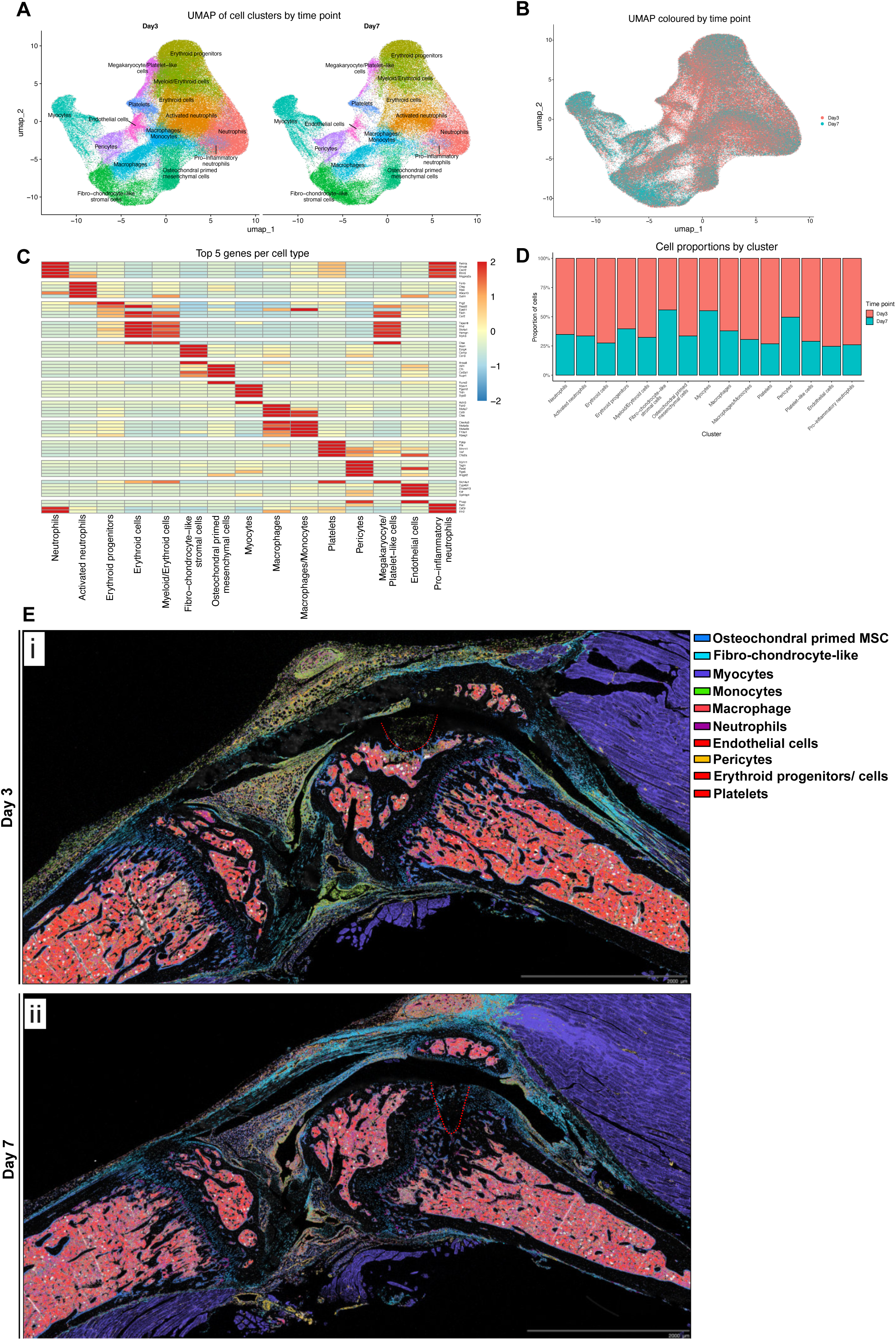
Spatial transcriptomic profiling revealing dynamic cellular composition across murine joint tissues at days 3 and 7 following injury. Spatial transcriptomics was performed on murine knee joints, 3 and 7 days following an osteochondral injury. (A) UMAP visualisation of all detected cell clusters at day 3 and day 7 post-injury, coloured by cell identity. (B) UMAP coloured by timepoint, illustrating the relative distribution of cells from day 3 and day 7 across clusters. (C) Heatmap showing the top 5 marker genes defining each identified cell population. (D) Proportional representation of cell clusters at each timepoint, highlighting shifts in cellular composition between day 3 and day 7. (E) Spatial mapping of annotated cell populations within the joint at day 3 (i) and day 7 (ii), demonstrating their localisation across synovial and injury-associated regions. n= 2. Scale bars: 2 mm.

To define the cellular landscape after injury, day 3 and day 7 datasets representing the entire joint were integrated following cell segmentation and transcript assignment. Unsupervised clustering and UMAP visualisation identified 14 transcriptionally distinct cell populations reflecting functional states phenotypically identifiable as immune, vascular, stromal and skeletal cells (Fig. 6A-C). The identification of these cell types was selected based on their top differentially expressed genes. Neutrophils comprised three transcriptional states: a homeostatic neutrophil population marked by selective *Retnla* and *Mmp8* expression; an activated neutrophil subset enriched for *Fcnb* and *Ctsg*; and a proinflammatory neutrophil state expressing *Retn*, *Csf3r* and *IL1r2*. Macrophage lineage populations were likewise heterogeneous, consisting of a reparative, anti-inflammatory macrophage state expressing *Folr2* and *Ms4a7*, and a transcriptional monocyte-macrophage population expressing *Ms4a6c* and *Ms4ab*, consistent with infiltration and maturation. Joint infiltrating blood derived cells were also detected within the joint space, including myeloid/erythroid cells (*Fech^low^, Slc4a1^low^, Rhd^high^*), erythroid progenitors (*Fech^Int^, Slc4a1^low^, Rhd^low^*) and more mature erythroid cells (*Fech^high^*, *Slc4a1^high^, Rhd^high^*), and platelet (Pf4, Ppbp, Vwf) and Megakaryocyte/platelet-like cells (Pf4, Ppbp and Slc4a1). Endothelial cells expressed Kdr and Plvap, while pericytes showed selective expression of *Myh11* and *Rgs5*, indicating vascular remodelling and a population of myocyte-like cells (*Myoz1, pgam2*) near the joint. Two skeletal lineage states were identified: osteochondral primed mesenchymal cells (OC-MSCs) marked by *Wif1* and *Cfh*, and fibro-chondrocyte-like stromal cells (FCSCs) expressing *Acan* and *Comp*. Collectively, these cell states reflect a dynamic spectrum of immune activation, hematopoietic contribution, and putative skeletal progenitor engagement that shape the early response to osteochondral injury.

To quantify temporal dynamics across the whole joint, cluster proportions were compared between the two timepoints (Fig. 6D). At day 3, the tissue environment was dominated by innate immune and haematopoietic populations, including neutrophils (across all three transcriptional states), monocytes, erythroid cells, erythroid progenitors, and platelet-related clusters, consistent with an acute inflammatory and haemostatic response. By day 7, these populations had markedly decreased, whilst macrophages comprised a greater proportion of cells, indicating a shift toward a macrophage-driven reparative phase. Endothelial cells and pericytes also declined over time, reflecting resolution of early vascular activation. Among the stromal populations, OC-MSCs showed a proportional decline by day 7, whereas FCSCs illustrated an increase over the same period, indicating a shift from early progenitor mobilisation toward expanding matrix-producing cell populations. Together, these changes illustrate a coordinated progression from acute immune infiltration and vascular activation to a macrophage-enriched and increasingly stromal repair environment during the first week of osteochondral repair.

Spatial mapping of transcriptionally distinct cell populations demonstrated region specific organisation across the joint (Fig. 6E). At day 3, neutrophil and monocyte-macrophage populations were concentrated within the appositional synovium and infiltrated the defect itself, consistent with early inflammatory recruitment to the site of tissue injury. Platelet-like and erythroid cells were aligned with the synovial and subchondral vascular regions, reflecting acute haemostatic activation. By day 7, monocyte and macrophage populations accumulated in the periarticular synovium, whereas OC-MSCs and FCSCs formed more organised clusters at the defect margins and base, marking the initiation of structural repair.

To identify differences in tissue distribution of cell types between day 3 and day 7 in the early repair response, we compared cell populations across four anatomically defined areas: (1) appositional synovium, (2) periarticular synovium, (3) defect surface, and (4) defect base (Fig. 7A-D). At day 3, OC-MSCs were preferentially localised to the defect base, whereas FCSCs were more prominent within the appositional synovium and were relatively limited within the defect itself (Fig. 7A). By day 7, FCSCs were increasingly represented at the defect surface and base, while OC-MSCs were less prominent, indicating a temporal shift in the composition and spatial organisation of the stromal repair response (Fig. 7B).

**Figure 7.**
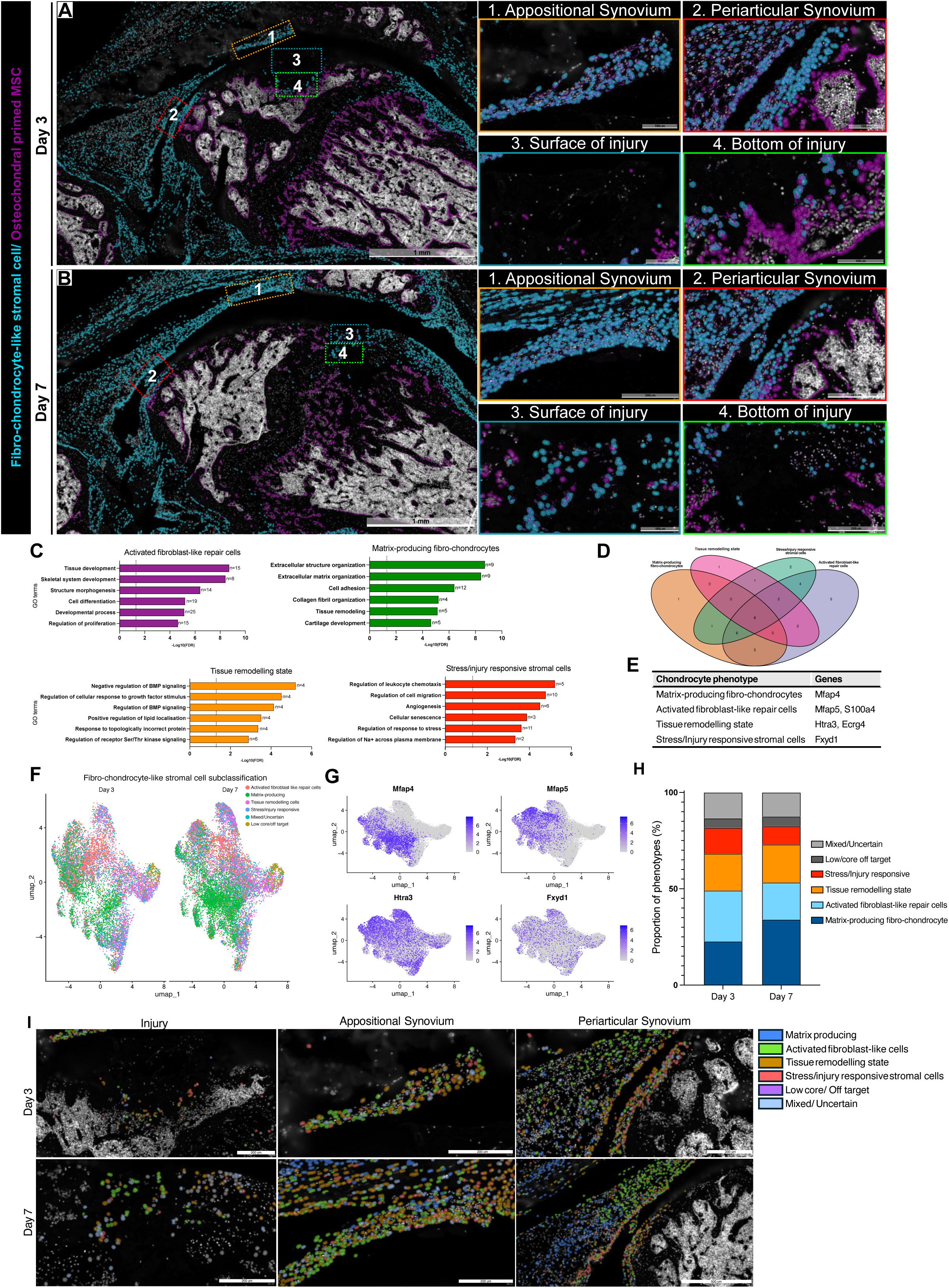
Functional heterogeneity and spatial organisation of fibro-chondrocyte-like stromal cell states during early osteochondral repair. (A-B) Spatial localisation of fibro-chondrocyte-like stromal cells and osteochondral primed MSCs at day 3 (A) and day 7 (B), with higher magnification views of defined anatomical regions including appositional synovium, periarticular synovium, injury surface, and injury base. (C) Gene ontology enrichment analysis of identified fibro-chondrocyte-like stromal subpopulations, including activated fibroblast-like repair cells, matrix-producing fibro- chondrocytes, tissue remodelling states, and stress/injury-responsive stromal cells. (D) Overlap of gene signatures across stromal phenotypes. (E) Summary of phenotype- associated anchor genes. (F) UMAP visualisation of fibro-chondrocyte-like stromal cell subclassification across timepoints. (G) Feature plots showing expression of representative marker genes (Mfap4, Mfap5, Htra3, Fxyd1) across stromal subpopulations. (H) Proportional distribution of stromal phenotypes at day 3 and day 7. (I) Spatial distribution of stromal subpopulations across the injury, appositional synovium, and periarticular synovium at day 3 and day 7. Data are presented as proportions or enrichment scores where appropriate. Scale bars: overview images = 1 mm; inset images = 200-500 µm.

### Region specific organisation of putative stromal populations during early osteochondral repair

To further examine the spatial organisation of skeletal lineage populations identified in Fig. 6, we analysed the localisation of OC-MSCs and FCSCs across distinct anatomical regions of the joint: (1) appositional synovium, (2) periarticular synovium, (3) surface of injury, and (4) base of the injury (Fig. 7A-B). Spatial distributions were compared between day 3 and day 7 post-injury.

At day 3 post-injury, OC-MSCs were predominantly localised at the base of the osteochondral injury, with a smaller population present along the injury surface (Fig. 7A). In contrast, FCSCs were present at lower abundance within the injury region at this time point. By day 7, both the surface and base of the injury showed increased presence of FCSCs, while OC-MSCs were restricted to the surface of the injury (Fig. 7B).

FCSCs were consistently detected within the appositional and periarticular synovium at both time points, with an increased abundance by day 7 (Fig. 7A1,2 and 7B1,2). OC-MSCs were undetected within the synovial regions at both time points.

### Fibro-chondrocyte-like stromal cells comprise distinct repair-associated transcriptional states

Given the spatial distribution of FCSCs across synovial and injury-associated regions, we next examined transcriptional heterogeneity within this population. Differential gene expression analysis followed by gene ontology enrichment identified four transcriptionally distinct FCSC phenotypes (Fig. 7C-E). These included: (1) matrix-associated fibro-chondrocytes, enriched for extracellular matrix organisation, cartilage development, and collagen fibril assembly pathways; (2) fibroblast-like stromal cells, associated with tissue development and regulation of protein secretion; (3) a tissue remodelling-associated state characterised by enrichment of Bone Morphogenic Protein (BMP) and growth factor signalling pathways; and (4) stress/injury-associated stromal cells enriched for inflammatory and stress-response pathways, including leukocyte chemotaxis and angiogenesis (Fig. 7C). The specificity of representative anchor genes for these phenotypes across timepoints is shown in Supplementary Fig. S1A, with quantitative marker performance metrics provided in Supplementary Fig. S1B.

Marker gene analysis supported these classifications (Fig. 7E). Matrix-associated fibro- chondrocytes were associated with extracellular matrix genes including *Mfap4*, while fibroblast-like stromal cells were marked by *Mfap5* and *S100a4*. Tissue remodelling-associated populations showed enrichment for genes including *Htra3* and *Ecrg4*, whereas stress/injury-associated stromal cells were characterised by expression of *Fxyd1*.

Projection of these gene signatures within the FCSC population confirmed the presence of transcriptionally distinct stromal sub-states occupying discrete regions of the cellular landscape (Fig. 7F-G). Quantification of their proportions across timepoints revealed temporal shifts in stromal state composition (Fig. 7H), with detailed cell counts and proportional representation summarised in Supplementary Fig. S1C. At day 3, fibroblast-like stromal cells and stress/injury-associated stromal cells were relatively enriched. By day 7, matrix-associated fibro-chondrocytes represented a larger proportion of the stromal compartment.

Spatial mapping of these stromal phenotypes revealed region-specific localisation patterns across the joint (Fig. 7I). Matrix-associated fibro-chondrocytes were largely absent from the injury region at both time points but were detected within synovial compartments, particularly by day 7. In contrast, activated fibroblast-like stromal cells and stress/injury-associated stromal cells were observed across both the injury region and synovial compartments. Tissue remodelling-associated stromal states were distributed across both synovial and injury-associated regions. Correlation analysis demonstrated that stress/injury-associated stromal cells were most closely related to activated fibroblast-like stromal cells (Supplementary Fig. S1D).

### Runx2-Sox9 lineage programmes delineate spatially distinct skeletal progenitor populations

To further define lineage-associated transcriptional states, we examined the expression of *Runx2* and *Sox9* across skeletal progenitor populations. Across all 14 transcriptionally defined cell populations (Fig. 6A-C), non-skeletal populations, including immune and vascular cell types, showed minimal expression of *Runx2* and *Sox9*. In contrast, FCSCs and OC-MSCs displayed distinct patterns of *Runx2* and *Sox9* positivity (Fig. 8A).

**Figure 8.**
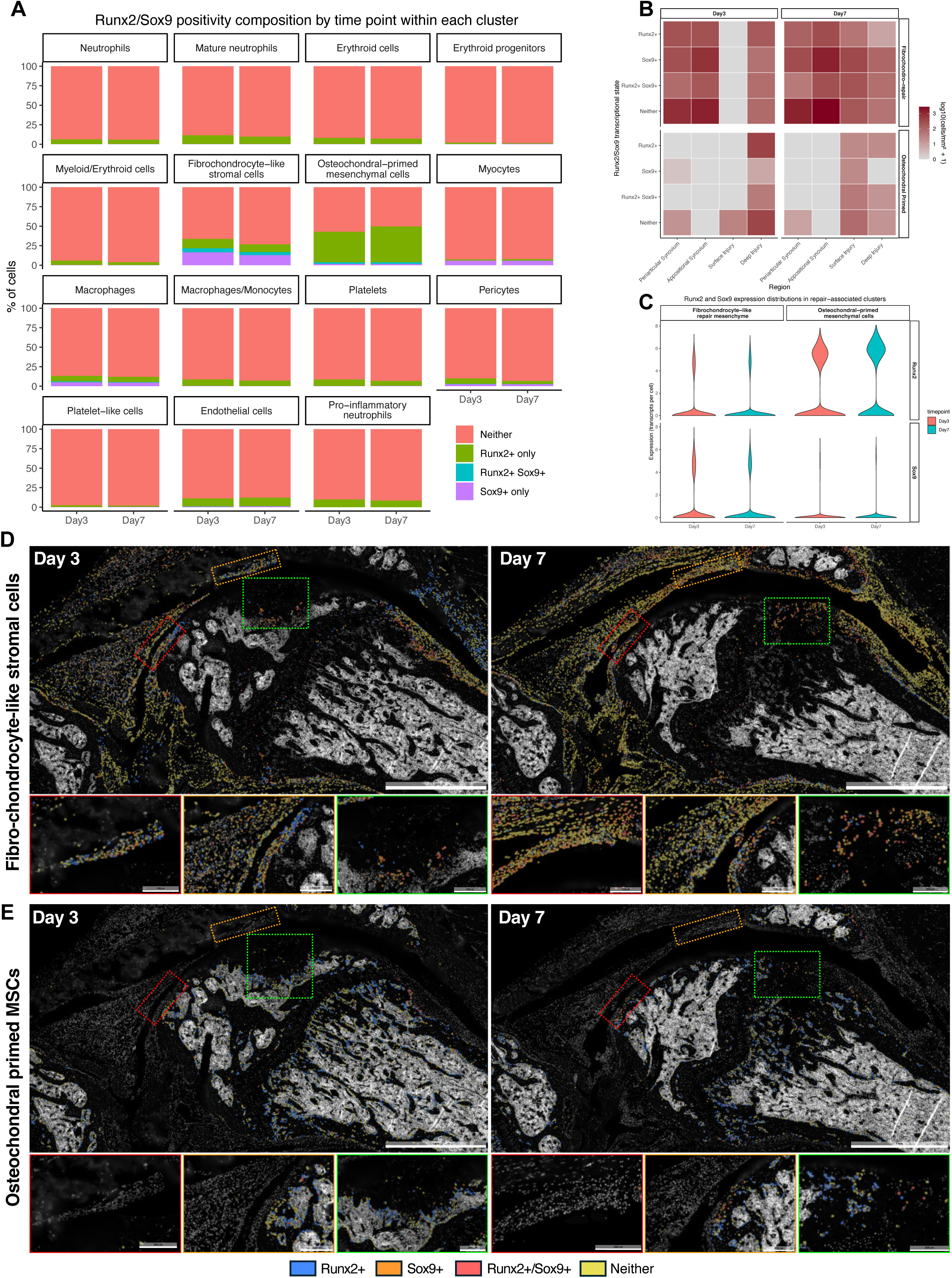
Spatial distribution of Runx2-Sox9 lineage-associated states across stromal populations during early osteochondral repair. (A) Composition of Runx2⁺, Sox9⁺, double-positive (Runx2⁺/Sox9⁺), and double-negative stromal cells across identified cell clusters at day 3 and day 7 post-injury. (B) Heatmap showing relative expression of lineage-associated genes across clusters and timepoints in different anatomical regions including the synovium and site of injury. (C) Distribution of Runx2 and Sox9 transcript levels within fibro-chondrocyte-like stromal cells and osteochondral primed MSCs at day 3 and day 7. (D) Spatial localisation of Runx2-Sox9 transcriptional states within fibro-chondrocyte-like stromal cells at day 3 and day 7, with higher magnification views of anatomically defined regions. (E) Spatial localisation of Runx2-Sox9 transcriptional states within osteochondral primed MSCs at day 3 and day 7. Colour coding indicates Runx2⁺ (blue), Sox9⁺ (orange), double-positive (red), and double-negative (yellow) cells. Data are presented as proportions or transcript counts where appropriate. Scale bars: overview images = 1 mm; inset images = 100-200 µm.

Within these populations, both FCSCs and OC-MSCs comprised mixtures of *Runx2*⁺, *Sox9*⁺, double-positive and double-negative cells. Double positive Runx2⁺/Sox9⁺ cells were predominantly observed within FCSCs and were present at lower frequency within OC-MSCs. Sox9⁺ cells were more abundant within FCSCs, whereas Runx2⁺ cells were more prevalent within OC-MSCs (Fig. 8A). Both populations exhibited a large proportion of double negative cells at both time points.

We next examined the spatial localisation of these transcriptional states across anatomically defined regions of the joint (Fig. 8B, D-E). Within FCSCs, all transcriptional states were distributed across both synovial and injury-associated regions at both time points. Sox9⁺ and double positive-Runx2⁺/Sox9⁺-cells were enriched within synovial compartments, particularly the appositional synovium, while Runx2-associated states were more frequently observed within injury-associated regions, predominantly at the base. By day 7, Sox9⁺ and dual-positive FCSCs were detected across both synovial and injury regions at increased proportions.

In contrast, OC-MSCs displayed a more regionally restricted distribution. At day 3, Runx2⁺ cells were predominantly localised to deeper regions of the injury, whereas Sox9 expression was less prominent. Dual Runx2⁺/Sox9⁺ cells were minimal at both time points. By day 7, OC- MSCs showed reduced representation across the whole injury, including decreased Runx2 expression and double positive cells deeper in the injury. These populations were, however, prevalent at the surface of the injury (Fig. 8B, D-E).

Quantitative analysis of gene expression further supported these observations (Fig. 8C). In FCSCs, both Runx2 and Sox9 were detected in subsets of cells at day 3 and day 7, with a higher proportion of Sox9⁺ cells. In contrast, OC-MSCs exhibited higher Runx2 expression overall, while Sox9 expression remained comparatively lower.

### Spatiotemporal organisation of immune cell infiltration during early osteochondral repair

We next examined the spatial distribution of immune cell populations across anatomically defined joint regions, including the appositional synovium, periarticular synovium, surface of the injury, and base of the injury (Fig. 9A-B).

**Figure 9.**
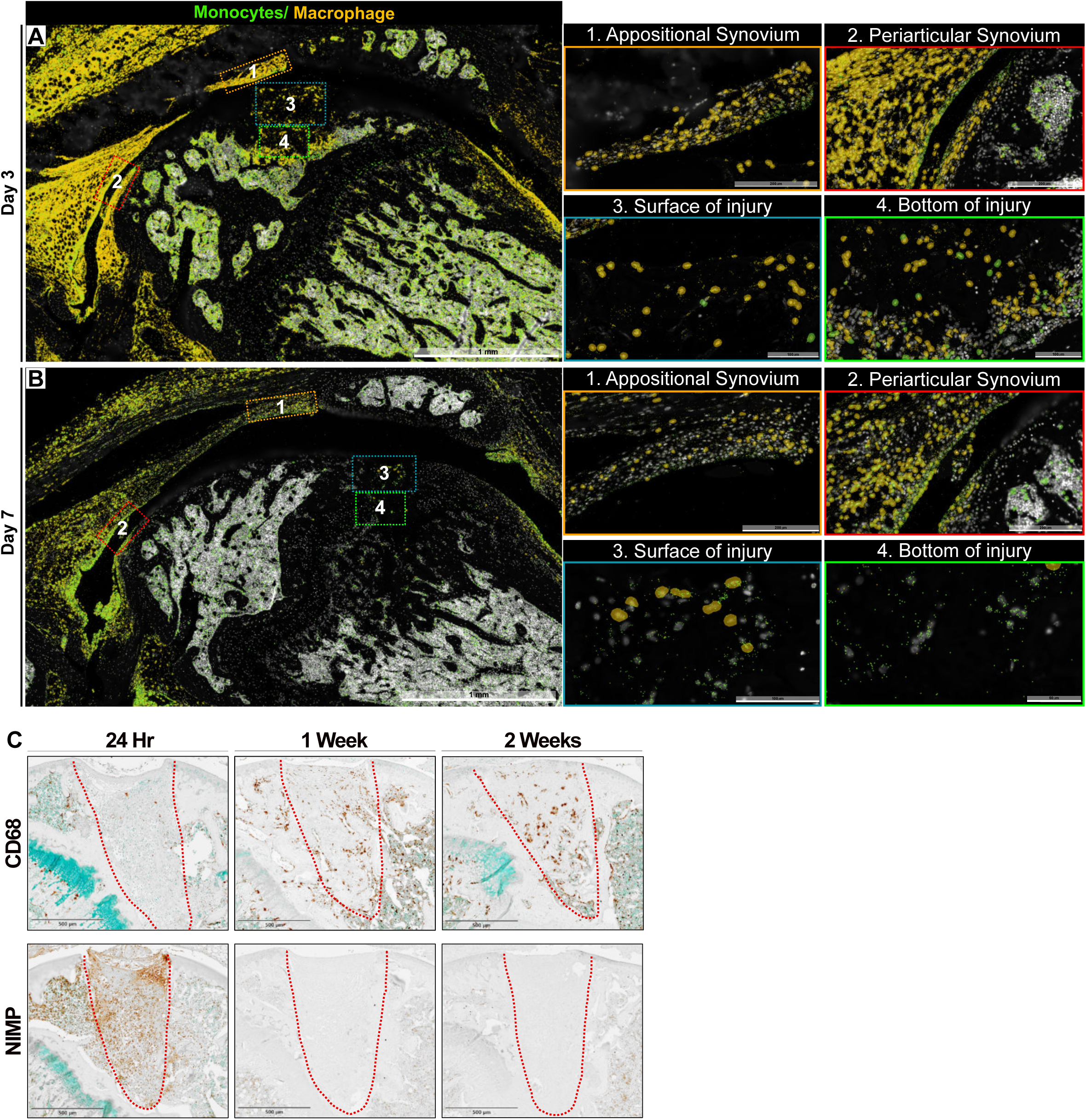
Spatial enrichment of immune cell infiltration within the synovium and injury during early osteochondral repair. (A-B) Spatial localisation of monocyte/macrophage populations within the joint at day 3 (A) and day 7 (B) post-injury, with higher magnification views of anatomically defined regions including appositional synovium, periarticular synovium, injury surface, and injury base. (C) Immunohistochemical staining for CD68⁺ macrophages and NIMP⁺ neutrophils within the defect at 24 hours, 1 week, and 2 weeks post-injury, illustrating early neutrophil presence followed by sustained macrophage localisation. Regions of interest within the defect are outlined with a red dashed line. Data are representative of n= 2 animals per timepoint. Scale bars: overview images = 1mm; inset images = 100-200 µm.

At day 3 post-injury, monocyte/macrophage populations were enriched within synovial compartments, particularly the appositional synovium, extending toward the site of injury (Fig. 9A). These cells were also detected throughout the injury. The periarticular synovium likewise demonstrated substantial immune cell presence.

By day 7, immune cell localisation was more spatially restricted (Fig. 9B). Monocyte/macrophage populations remained detectable within synovial regions albeit fewer in number but were reduced within deeper regions of the injury, with a continuous presence along the surface of the injury.

To further characterise these dynamics, immunohistochemical analysis was performed using NIMP (neutrophils) and CD68 (macrophages) at 24 hours, 1 week, and 2 weeks post-injury (Fig. 9C). At 24 hours, NIMP⁺ neutrophils were abundantly present within the defect. By 1 week, neutrophils were largely absent and remained minimal at 2 weeks. In contrast, CD68⁺ macrophages were detected within both the surface and deeper regions of the defect from 1 week and remained present at 2 weeks.

## Discussion

This study defines the early cellular organisation of osteochondral repair by using a reproducible murine injury model and integrating spatial transcriptomics, immunophenotyping, histology and longitudinal imaging data. To our knowledge, it provides the first spatially resolved transcriptomic analysis of the early response to focal osteochondral injury across the intact synovial joint. The principal finding is that the early repair response is established through the emergence of anatomically distinct immune, vascular and stromal microenvironments during the first week after injury. An initial inflammatory and haemostatic response gave way to increasingly organised cellular niches, while transcriptionally distinct stromal populations occupied discrete synovial and injury-associated locations. Histology, MRI and µCT extended these observations, demonstrating progressive but incomplete restoration of osteochondral architecture, transient synovial and vascular activation, and more effective regeneration of subchondral bone than articular cartilage. Together, these findings identify the first week after injury as a critical phase during which the repair microenvironment is established, defining a biological framework from which subsequent osteochondral repair emerges.

### Spatial organisation defines the early repair response

At day 3 after injury, the joint is dominated by neutrophils, monocytes, erythroid cells and platelet-related populations distributed across both the defect and adjacent synovium. This profile is consistent with marrow disruption, clot formation and acute inflammatory cell recruitment following osteochondral injury (Shapiro et al., 1993). By day 7, these populations had declined, and macrophages now represent a greater proportion of the cellular environment. Immunohistochemistry supported this view of the temporal transition: neutrophils abundant within the defect at 24 hours but largely absent by one week, while macrophages persisting within the superficial and deeper repair regions. These findings indicate a rapid shift from acute inflammatory and haemostatic activity towards a macrophage-enriched repair environment.

The spatial data add an important dimension to this established sequence. At day 3, monocyte and macrophage populations were broadly distributed across the synovium and defect. By day 7, their localisation became more restricted, with continued representation along the injury surface and within adjacent synovial regions. This reorganisation coincided with the emergence of increasingly distinct stromal niches. Rather than demonstrating a simple replacement of immune cells by reparative cells, the data suggest that immune and stromal responses become anatomically organised within the injured joint. Such localisation may be functionally important because macrophages can regulate stromal-cell activation, matrix remodelling and differentiation through context-dependent inflammatory and reparative signals (Wynn & Vannella, 2016; Vi et al., 2015; Wu et al., 2020).

Two transcriptionally defined stromal populations contributed prominently to this organisation: fibro-chondrocyte-like stromal cells (FCSCs) and osteochondral-primed mesenchymal cells (OC-MSCs). OC-MSCs, marked by *Wif1* and *Cfh*, were most prominent during the earlier response and preferentially associated with the injury, particularly its deeper regions. Their relative abundance declined by day 7. In contrast, *Acan*/*Comp*-expressing FCSCs increased over the same period and occupied both synovial and injury-associated regions. This temporal shift is consistent with progression from an early osteochondral-associated stromal state towards a repair environment increasingly populated by fibro-chondrocyte-like cells. However, these transcriptional identities do not establish developmental origin or fate, and the designation of OC-MSCs should therefore be interpreted as describing a molecular state rather than a proven progenitor lineage.

The regional distribution of these populations suggests that the injury surface, base and synovium provide distinct cellular environments. OC-MSCs were preferentially associated with the osteochondral injury, whereas FCSCs were prominent within the synovium and became increasingly represented within the repair tissue by day 7. The presence of related FCSC populations in the synovium and defect raises the possibility that the synovium contributes to the repair-associated stromal environment. This interpretation is consistent with previous evidence that synovial stromal cells possess chondrogenic and extracellular matrix-producing capacity (De Bari et al., 2001; Sakaguchi et al., 2005; Roelofs et al., 2017; Zamudio-Cuevas et al., 2022). Nevertheless, spatial proximity cannot distinguish migration from local activation, convergent transcriptional states or independent resident populations. The synovium can therefore be considered a potential stromal niche or reservoir, but its direct cellular contribution to the defect requires lineage tracing.

### Regional anatomical niches support distinct stromal states

FCSCs were not transcriptionally uniform. They comprised matrix-associated, fibroblast-like, tissue-remodelling and stress/injury-associated states whose representation changed with time and location. Fibroblast-like and stress-responsive states were more prominent during the early response, whereas matrix-associated FCSCs increased by day 7. This transition indicates that stromal activation is dynamic and supports the broader concept that fibroblast identity is shaped by tissue context rather than representing a fixed cell type (Muhl et al., 2020; Zhang et al., 2019; Buechler et al., 2021).

The anatomical distribution of these states was particularly informative. Matrix-associated FCSCs were preferentially detected within synovial compartments, while the defect remained enriched for fibroblast-like, remodelling and stress-responsive states. The injury environment therefore contained stromal cells that did not exhibit the same matrix-associated programme observed within the synovium during this early period. This distinction suggests that incomplete cartilage regeneration may reflect not only the availability of repair-responsive cells, but also whether the local environment permits them to establish and sustain an appropriate matrix-forming phenotype.

The persistence of activated and stress-responsive states within the defect is compatible with continued exposure to inflammatory, mechanical and remodelling signals. In contrast, preservation of tissue architecture and a different signalling environment within the synovium may favour matrix-associated programmes. The greater representation of these states in synovial regions more distant from the defect further suggests a spatial gradient in injury-associated cues. Although the targeted gene panel cannot define the responsible signalling pathways comprehensively, these findings identify the local microenvironment as a plausible determinant of stromal function during early repair.

Analysis of *Runx2* and *Sox9* expression further distinguished the stromal populations. FCSCs contained a greater proportion of *Sox9*-positive and *Runx2*/*Sox9* double-positive cells, particularly within synovial regions, consistent with a fibro-chondrogenic transcriptional programme. *Sox9* is central to chondrogenic identity and cartilage extracellular matrix regulation, supporting the association between these cells and matrix-producing potential (Lefebvre & Dvir-Ginzberg, 2017). OC-MSCs showed comparatively greater *Runx2* expression and were concentrated within injury-associated regions, particularly the base of the osteochondral injury. *Runx2* regulates osteogenic commitment and hypertrophic progression during skeletal development and repair, making this profile compatible with an osteochondral-associated state (Komori, 2020).

The presence of *Runx2*/*Sox9* double-positive cells, most prominently among FCSCs, may represent transitional transcriptional states rather than established lineages. Co-expression of these factors has been reported during skeletal development and repair, where it can precede resolution towards osteogenic or chondrogenic programmes (Zhou et al., 2015; Tsang & Cheah, 2019; Wang et al., 2022). Their spatial enrichment within synovial and injury-associated niches suggests that lineage-associated programmes remain plastic during the first week after injury. However, many FCSCs and OC-MSCs lacked detectable *Runx2* or *Sox9* expression, indicating that these repair-associated stromal populations are defined by broader transcriptional states rather than expression of individual lineage markers alone. This observation highlights the molecular heterogeneity of stromal cells during early repair, although the limited transcript panel and sensitivity of targeted spatial profiling should be considered when interpreting marker-negative populations.

The data therefore support a continuum of stromal activation and differentiation states rather than a binary separation into committed skeletal cell types.

### Early spatial organisation is associated with the structural trajectory of repair

The later structural findings were consistent with the early organisation revealed by spatial transcriptomics. Histologically, the defect initially contained a provisional erythrocyte-rich clot, followed by a highly cellular repair tissue and focal proteoglycan deposition. From two weeks onwards, repair became anatomically organised: proteoglycan-containing matrix developed towards the articular surface, while mineralised tissue regenerated from the base of the injury. This superficial-to-deep separation mirrors the early distinction between fibro-chondrocyte-like stromal states and Runx2-enriched osteochondral-associated cells.

Longitudinal µCT demonstrated rapid regeneration of the osseous compartment. Initial loss of subchondral bone was followed by peripheral bridging and progressive mineralised infilling between weeks 1 and 8. This sequence resembles the response to marrow-stimulation procedures, in which subchondral bone repair can precede maturation of the overlying cartilage (Kreuz et al., 2006; Solheim et al., 2016). Despite substantial infilling, trabecular architecture remained irregular at eight weeks, indicating continued remodelling. Histology likewise showed re-establishment of an osteochondral boundary without complete restoration of native architecture. These findings demonstrate that successful osseous repair does not necessarily restore the structure or composition of the overlying articular cartilage.

The composition of the late repair tissue further supports this conclusion. COL1A1 was prominent within the surface region, whereas COL2A1 was restricted and discontinuous. COL10A1 persisted around the osteochondral interface and extended into the repair tissue, while PRG4 was restored at the articular surface. The regenerated tissue therefore acquired some features of an osteochondral surface, including proteoglycan deposition and a lubricating boundary layer, but retained fibrous and hypertrophic characteristics. This phenotype is consistent with the fibrocartilaginous repair produced after marrow stimulation and helps explain why structural filling does not necessarily confer durable hyaline cartilage restoration (Mithoefer et al., 2009; Minas & Ogura, 2016; Williams et al., 2019; Solheim et al., 2020).

These findings several weeks after injury provide important context for the spatial transcriptomic observations. The early defect is not devoid of stromal cells or matrix-associated potential. Rather, matrix-associated FCSC states appear preferentially supported within synovial compartments, while activated and stress-responsive states persist within the injury. One interpretation is that the local defect environment does not adequately stabilise a hyaline cartilage-producing programme during the critical early phase. The eventual formation of COL1A1-rich, COL2A1-limited repair tissue may therefore reflect an early mismatch between stromal potential and the signals operating within the injury niche. This relationship remains associative, but it provides a testable framework for future intervention.

### Regional synovial and vascular niches define the early repair environment

Spatial transcriptomics also identified early endothelial and pericyte populations that declined proportionally between days 3 and 7, indicating rapid modification of the vascular environment. MRI and histomorphometry extended this observation beyond the first week. Synovial thickening and MRI hyperintensity were evident early, increased towards week 4 and partially resolved by week 8. Vessel area fraction declined across the study, whereas vessel size showed a transient increase, particularly within the appositional synovium. These complementary measurements indicate that vascular remodelling involved both changes in vascular representation and temporary enlargement of individual vessels.

The distinction between appositional and periarticular synovium was also biologically informative. The appositional synovium, positioned directly against the defect, remained thicker and showed more pronounced vascular changes than the periarticular lining. This regional response supports the spatial transcriptomic finding that the synovium is not a uniform compartment. Proximity to the injury appears to shape immune, vascular and stromal behaviour, producing local environments with potentially different contributions to repair.

Synovitis and vascular remodelling are increasingly recognised as active components of joint injury rather than secondary consequences of cartilage damage (Atukorala et al., 2016; Sanchez-Lopez et al., 2022; Knights et al., 2023). Early vascular activation may facilitate immune-cell entry, nutrient delivery and tissue remodelling. However, persistent inflammatory and pro-angiogenic signalling can also promote fibrosis, hypertrophy and osteogenic differentiation, processes incompatible with stable hyaline cartilage. In this model, synovial and vascular activation partially resolved, yet the final cartilage remained fibrocartilaginous and hypertrophic. It cannot be concluded that the vascular response caused this outcome, but its timing overlaps the period during which stromal states were being established. The synovial-vascular environment may therefore help determine whether repair-associated stromal cells adopt stable chondrogenic, fibrotic or hypertrophic programmes.

### Implications of osteochondral repair and regenerative therapy

The focal injury model complements, rather than replaces, instability-based models such as destabilisation of the medial meniscus. DMM reproduces progressive joint-wide tissue degeneration driven by altered mechanics and chronic inflammation (Glasson et al., 2007), whereas the injury presented here isolates the acute response to a defined osteochondral lesion. Controlled penetration of the marrow compartment reproduces a central feature of marrow-stimulation procedures by allowing blood, marrow-derived cells and inflammatory mediators to access the defect (Shapiro et al., 1993; Frisbie et al., 2006; Erggelet & Vavken, 2016). Its reproducible geometry also permits consistent regional analysis and repeated MRI and µCT assessment within the same animal.

This model is therefore suited to studying the early regenerative-degenerative balance following focal injury. Localised osteochondral defects are clinically important because they affect younger patients and increase the risk of post-traumatic osteoarthritis. Current marrow- stimulation procedures can produce defect filling but commonly generate mechanically inferior fibrocartilage that deteriorates with time (Mithoefer et al., 2009; Lieberthal et al., 2015; Solheim et al., 2020). The present findings suggest that improving repair may require more than supplying additional cells or stimulating matrix production. Effective intervention may need to address the immune, vascular and stromal environments during the period when spatial repair niches are first established.

These findings suggest that successful osteochondral repair may depend on spatially and temporally targeted modulation of the repair microenvironment, rather than interventions directed at individual tissues or cell populations alone. Early inflammatory responses are likely to require modulation rather than suppression, as they initiate repair and facilitate debris clearance. Similarly, macrophage activity may require temporal and regional control to preserve reparative functions while limiting persistent inflammatory or fibrotic signalling. Stromal-directed therapies may need to promote matrix-associated cellular states specifically within the defect, where these programmes were underrepresented, rather than simply increasing stromal-cell abundance throughout the joint. Finally, strategies that coordinate regeneration of articular cartilage with remodelling of the underlying subchondral bone may prove more effective than approaches targeting either compartment in isolation. Together, these findings establish a framework for developing and evaluating therapies that are directed towards the evolving repair microenvironment in both space and time.

### Limitations and future directions

The spatial framework presented here defines several priorities for future investigation. Developing a more comprehensive spatial atlas of the critical early repair window, incorporating larger cohorts, additional time points and expanded gene panels, will enable finer resolution of the cellular states and signalling networks that coordinate osteochondral repair. Accordingly, the populations identified in this study should be interpreted as transcriptionally defined repair states rather than definitive lineage classifications. Although the present analyses reveal distinct spatial organisation across the injured joint, they do not establish developmental origin, migration or functional contribution. Integrating genetic lineage tracing, fate mapping and functional manipulation will be essential to determine how these repair-associated populations contribute to osteochondral regeneration and how they may be targeted therapeutically. Furthermore, the present study was performed exclusively in young adult female mice, and future studies should determine whether these spatial programmes are conserved across sex, age and other models of osteochondral injury.

## Conclusion

In conclusion, this study identifies the progressive spatial organisation of the early osteochondral repair response as a defining feature of joint healing following osteochondral injury. Acute immune and haemostatic activity gives way to a macrophage-enriched repair environment, while transcriptionally distinct stromal populations become regionally organised across the synovium, defect surface and defect base. Matrix-associated fibro-chondrocyte-like states are enriched within the synovium, whereas activated and stress-responsive stromal programmes predominate within the injury site during the first week after injury. Longitudinal histological, MRI and µCT analyses demonstrate that these early biological events are followed by progressive restoration of osteochondral architecture, with more effective regeneration of subchondral bone than native articular cartilage, resulting in structurally integrated but compositionally immature repair tissue. Together, these findings suggest that the limited regenerative capacity of articular cartilage reflects not simply a shortage of reparative cells, but an inability to establish and sustain the coordinated immune, vascular and stromal microenvironment required for successful regeneration. By integrating spatial transcriptomic profiling with longitudinal structural assessment, this study establishes a biological framework for understanding early osteochondral repair and provides a foundation for the rational design of spatially and temporally targeted regenerative therapies.

## Abbreviations

ACI: Autologous Chondrocyte Implantation
BMP: Bone Morphogenic Protein
BSA: Bovine Serum Albumin
DMM: Destabilisation of the Medial Meniscus
EDTA: Ethylenediaminetetraacetic acid
FCSC: Fibro-chondrocyte-like stromal cells
GO: Gene Ontology
LSD: Fisher’s Least Significant Difference
MRI: Magnetic Resonance Imaging
MSME: Multi-Slice Multi-Echo
OC-MSC: Osteochondral-primed mesenchymal stromal cells
PBS: Phosphate Buffered Saline
PC: Principal Components
PCA: Principal Component Analysis
PTOA: Post-traumatic osteoarthritis
RARE: Rapid Acquisition with Refocusing Echoes
ROI: Regions of Interest
SD: Standard Deviation
TBS: Tris Buffered Saline
UMAP: Uniform Manifold Approximation and Projection

---

**Supplementary Figure 1. Phenotypic classification and gene signature validation of fibro-chondrocyte-like stromal cells (FCSCs)** (A) Dot plot of anchor gene expression across FCSC phenotypes at Day 3 and Day 7. Dot size indicates the percentage of expressing cells and colour represents average expression, demonstrating phenotype-specific transcriptional profiles. (B) Summary of anchor gene performance for phenotype classification, including target association, AUC, log fold change, detection rates, and specificity ratio. (C) Quantification of FCSC phenotypes at Day 3 and Day 7, showing cell numbers and proportions across fibroblast-like repair, matrix-producing, tissue remodelling, stress/injury-responsive, mixed/uncertain, and low core/off-target states. (D) Correlation heatmap of stress/injury-responsive cells with other FCSC phenotypes, showing transcriptional relationships between stromal states.

## Acknowledgements

The authors gratefully acknowledge Dr Francesca M. D. Henson for establishing the original Home Office Project Licence and for providing technical guidance and continued support during the development and initial validation of the murine osteochondral injury model. The authors also thank Dr Karin Newell for her technical expertise and support with the animal studies and histological processing.

## Funders

This work was supported by the Medical Research Council through the UK Regenerative Medicine Platform and by the ALBORADA Trust.

## Notes

### Competing Interest Statement

The authors have declared no competing interest.

## References

Armiento, A. R., Alini, M. & Stoddart, M. J. Articular fibrocartilage- Why does hyaline cartilage fail to repair? Adv. Drug Deliv. Rev. 146, 289–305 (2019).

Atukorala, I., Kwoh, C. K., Guermazi, A., Roemer, F. W., Boudreau, R. M., & Hunter, D. J. Synovitis in knee osteoarthritis: a precursor of disease? Ann. Rheum. Dis. 75, 390–395 (2016).

Buechler, M. B. et al. Cross-tissue organization of the fibroblast lineage. Nature 593, 575–579 (2021).

Curl, W. W. et al. Cartilage injuries: a review of 31,516 knee arthroscopies. Arthroscopy 13, 456–460 (1997).

De Bari, C., Dell’Accio, F. & Luyten, F. P. Human periosteum-derived cells maintain phenotypic stability and chondrogenic potential throughout expansion regardless of donor age. Arthritis Rheum. 44, 85–95 (2001).

Eckstein, F., Collins, J. E., Nevitt, M. C., Lynch, J. A., Kraus, V. B. & Katz, J. N. Cartilage thickness change as an imaging biomarker of knee osteoarthritis progression. Nat. Rev. Rheumatol. 16, 343–357 (2020).

Eltawil NM, De Bari C, Achan P, Pitzalis C, Dell’accio F. A novel in vivo murine model of cartilage regeneration. Age and strain-dependent outcome after joint surface injury. Osteoarthritis Cartilage. Jun;17(6):695–704 (2009).

Erggelet, C. & Vavken, P. Microfracture for the treatment of cartilage defects in the knee joint - A golden standard? J. Clin. Orthop. Trauma 7, 145–152 (2016).

Fitzgerald J, Rich C, Burkhardt D, Allen J, Herzka AS, Little CB. Evidence for articular cartilage regeneration in MRL/MpJ mice. Osteoarthritis Cartilage. (2008) Nov;16(11):1319–26.

Frisbie, D. D. et al. Early events in cartilage repair after subchondral bone microfracture. Clin. Orthop. Relat. Res. 442, 215–227 (2006).

Glasson, S. S., Blanchet, T. J. & Morris, E. A. The surgical destabilization of the medial meniscus (DMM) model of osteoarthritis in the 129/SvEv mouse. Osteoarthritis Cartilage 15, 1061–1069 (2007).

Gracitelli, G. C. et al. Surgical interventions for articular cartilage defects in the knee. J. Bone Joint Surg. Am. 97, 132–142 (2015).

Hunziker, E. B. Articular cartilage repair: basic science and clinical progress. A review of the current status and prospects. Osteoarthritis Cartilage 10, 432–463 (2002).

Knights, A. J. et al. Synovial tissue responses to joint injury: linking inflammation, vascularity and repair. Nat. Rev. Rheumatol. 19, 123–138 (2023).

Komori, T. Runx2, an inducer of osteoblast and chondrocyte differentiation. Histochem. Cell Biol. 154, 325–339 (2020).

Kreuz, P. C. et al. Results after microfracture of full-thickness chondral defects in different compartments in the knee. Osteoarthritis Cartilage 14, 1119–1125 (2006).

Kuroda, R. et al. Cartilage repair using bone morphogenetic protein 4 and muscle-derived stem cells. Arthritis Rheum. 54, 433–442 (2006).

Lieberthal, J., Sambamurthy, N. & Scanzello, C. R. Inflammation in joint injury and post-traumatic osteoarthritis. Osteoarthritis Cartilage 23, 1825–1834 (2015).

Little, C. B. & Hunter, D. J. Post-traumatic osteoarthritis: from mouse models to clinical trials. Nat. Rev. Rheumatol. 9, 485–497 (2013).

Luyten, F. P., Denti, M., Filardo, G., Kon, E. & Engebretsen, L. Definition and classification of early osteoarthritis of the knee. Knee Surg. Sports Traumatol. Arthrosc. 20, 401–406 (2012).

Madry, H. et al. Early osteoarthritis of the knee. Knee Surg. Sports Traumatol. Arthrosc. 24, 1753–1762 (2016).

Mahmoudian, A. et al. Early-stage symptomatic osteoarthritis of the knee—time for action. Nat. Rev. Rheumatol. 17, 621–632 (2021).

Matsuoka M, Onodera T, Sasazawa F, Momma D, Baba R, Hontani K, Iwasaki N. An Articular Cartilage Repair Model in Common C57Bl/6 Mice. Tissue Eng Part C Methods. (2015) Aug;21(8):767–72.

Minas, T. & Ogura, T, Bryant T. Autologous chondrocyte implantation. JBJS Essent Surg Tech 6(2):e24. (2016).

Mithoefer, K., McAdams, T., Williams, R. J., Kreuz, P. C. & Mandelbaum, B. R. Clinical efficacy of the microfracture technique for articular cartilage repair in the knee. Am. J. Sports Med. 37, 2053–2063 (2009).

Muhl, L. et al. Single-cell analysis uncovers fibroblast heterogeneity and criteria for fibroblast and mural cell identification. Nat. Commun. 11, 3953 (2020).

Pridie, K. H. A method of resurfacing osteoarthritic knee joints. J. Bone Joint Surg. Br. 41-B, 618–619 (1959).

Roelofs, A. J. et al. Identification of resident fibroblast progenitor cells in the synovial joint. Nat. Commun. 8, 15076 (2017).

Sakaguchi, Y. et al. Comparison of human stem cells derived from various mesenchymal tissues. Arthritis Rheum. 52, 2521–2529 (2005).

Sanchez-Lopez, E. et al. Synovial inflammation in osteoarthritis progression. Nat. Rev. Rheumatol. 18, 258–275 (2022).

Shapiro, F., Koide, S. & Glimcher, M. J. Cell origin and differentiation in the repair of full-thickness defects of articular cartilage. J. Bone Joint Surg. Am. 75, 532–553 (1993).

Solheim, E. et al. Long-term results after microfracture treatment of articular cartilage defects in the knee. Knee Surg. Sports Traumatol. Arthrosc. 24, 1587–1593 (2016).

Solheim, E. et al. Results at 10-14 years after microfracture treatment of articular cartilage defects in the knee. Knee Surg. Sports Traumatol. Arthrosc. 28, 2110–2117 (2016).

Solheim, E. et al. Long-term clinical outcomes after cartilage repair. Knee Surg. Sports Traumatol. Arthrosc. 28, 1466–1473 (2020).

Tsang, K. Y. & Cheah, K. S. E. The extended roles of Sox9 in cartilage and skeletal development. Curr. Osteoporos. Rep. 17, 213–219 (2019).

van der Kraan, P. M. The changing role of TGFβ in healthy, ageing and osteoarthritic joints. Nat. Rev. Rheumatol. 13, 155–163 (2017).

Versus Arthritis. Osteoarthritis in the UK. Available at: https://www.versusarthritis.org (accessed 2026).

Wang, Y. et al. Single-cell transcriptomics of skeletal development and repair. Nat. Commun. 13, 1234 (2022).

Williams, R. J. et al. Long-term outcomes of cartilage repair. J. Bone Joint Surg. Am. 101, 1029–1038 (2019).

World Health Organization. Osteoarthritis. Available at: https://www.who.int/news-room/fact-sheets/detail/osteoarthritis (accessed 2026).

Wu, C. L. et al. Macrophage biology in osteoarthritis. Nat. Rev. Rheumatol. 16, 123–137 (2020).

Wynn, T. A. & Vannella, K. M. Macrophages in tissue repair, regeneration, and fibrosis. Immunity 44, 450–462 (2016).

Zamudio-Cuevas, Y. et al. Synovial fibroblast heterogeneity in joint disease. Front. Immunol. 13, 845321 (2022).

Zhang, F. et al. Defining inflammatory cell states in rheumatoid arthritis synovial tissues. Nat. Immunol. 20, 928–942 (2019).

Zhou, X. et al. Chondrocytes transdifferentiate into osteoblasts in endochondral bone formation. Cell Metab. 21, 713–724 (2015).

