## Supplementary Figure 1 for "Spatiotemporal transcriptomic landscape of synovial joint repair – an in vivo murine multimodal model of osteochondral injury"

A

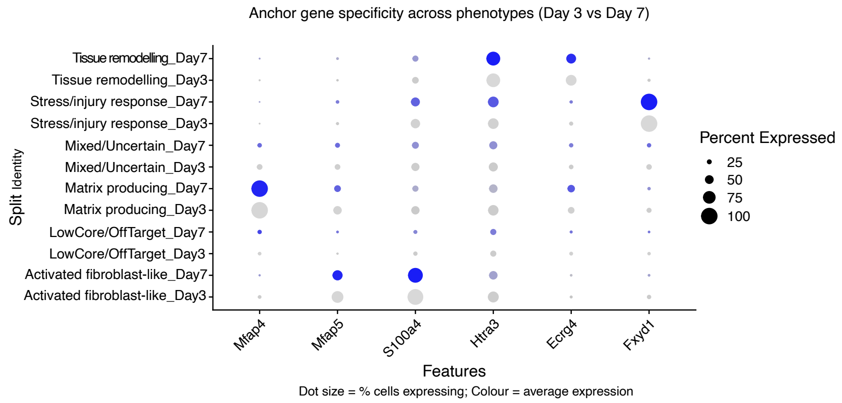


B

| Time point | Gene | Target Phenotype | No. target | No. other | AUC | logFC | cohens_d | pct_detect_target | pct_detect_other | specificity_ratio |
|---|---|---|---|---|---|---|---|---|---|---|
| Day3 | Mfap4 | Matrix producing Fibro-chondrocytes | 2762 | 9413 | 0.98 | 5.08 | 3.37 | 1.00 | 0.15 | 18.59 |
| | Mfap5 | Activated fibroblast-like | 3222 | 8953 | 0.73 | 2.25 | 0.99 | 0.69 | 0.26 | 2.98 |
| | S100a4 | Activated fibroblast-like | 3222 | 8953 | 0.83 | 3.09 | 1.32 | 0.95 | 0.44 | 3.35 |
| | Htra3 | Tissue remodelling | 2330 | 9845 | 0.67 | 1.44 | 0.57 | 0.82 | 0.59 | 1.82 |
| | Ecrg4 | Tissue remodelling | 2330 | 9845 | 0.72 | 2.25 | 1.03 | 0.63 | 0.23 | 4.40 |
| | Fxyd1 | Stress/Injury responsive | 1621 | 10554 | 0.94 | 3.99 | 2.32 | 1.00 | 0.22 | 7.95 |
| Day7 | Mfap4 | Matrix producing Fibro-chondrocytes | 5242 | 10159 | 0.98 | 5.59 | 4.19 | 1.00 | 0.10 | 24.37 |
| | Mfap5 | Activated fibroblast-like | 2966 | 12435 | 0.69 | 1.90 | 0.82 | 0.59 | 0.25 | 3.19 |
| | S100a4 | Activated fibroblast-like | 2966 | 12435 | 0.83 | 3.21 | 1.35 | 0.89 | 0.35 | 4.06 |
| | Htra3 | Tissue remodelling | 3035 | 12366 | 0.72 | 2.02 | 0.79 | 0.83 | 0.49 | 2.24 |
| | Ecrg4 | Tissue remodelling | 3035 | 12366 | 0.67 | 1.75 | 0.75 | 0.57 | 0.26 | 3.32 |
| | Fxyd1 | Stress/Injury responsive | 1470 | 13931 | 0.96 | 4.38 | 2.89 | 1.00 | 0.13 | 10.62 |

C

| Day 3 | | |
|---|---|---|
| **Phenotype** | **No of Cells** | **Proportion (%)** |
| Fibroblast-like repair cells | 3222 | 26.5 |
| Matrix-producing fibro-chondrocyte | 2762 | 22.7 |
| Tissue remodelling cells | 2330 | 19.1 |
| Stress/Injury responsive cells | 1621 | 13.3 |
| Mixed/Uncertain | 1631 | 13.4 |
| Low Core/Off Target | 609 | 5.0 |

| Day 7 | | |
|---|---|---|
| **Phenotype** | **No of Cells** | **Proportion (%)** |
| Fibroblast-like repair cells | 2966 | 19.3 |
| Matrix-producing fibro-chondrocyte | 5242 | 34.0 |
| Tissue remodelling cells | 3035 | 19.7 |
| Stress/Injury responsive cells | 1470 | 9.5 |
| Mixed/Uncertain | 1919 | 12.5 |
| Low Core/Off Target | 769 | 5.0 |

D

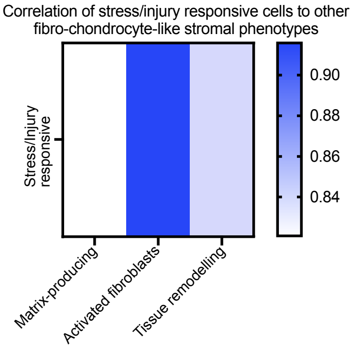


**Supplementary Figure 1. Phenotypic classification and gene signature validation of fibro-chondrocyte-like stromal cells (FCSCs)**
